# Improved reference genome uncovers novel sex-linked regions in the guppy (*Poecilia reticulata*)

**DOI:** 10.1101/2020.08.07.240986

**Authors:** Bonnie A. Fraser, James R. Whiting, Josephine R. Paris, Cameron J. Weadick, Paul J. Parsons, Deborah Charlesworth, Roberta Bergero, Felix Bemm, Margarete Hoffmann, Verena A. Kottler, Chang Liu, Christine Dreyer, Detlef Weigel

**Affiliations:** Biosciences, University of Exeter, Stocker Road, Exeter, EX4 4QD UK; Institute of Evolutionary Biology, School of Biological Sciences, University of Edinburgh, Edinburgh, EH9 3LF, UK; Department of Molecular Biology, Max Planck Institute for Developmental Biology, Max-Plank-Ring 5, 72076 Tübingen, Germany; Institute of Biology, University of Hohenheim, Garbenstraße 30, 70599 Stuttgart, Germany

## Abstract

Theory predicts that the sexes can achieve greater fitness if loci with sexually antagonistic polymorphisms become linked to the sex determining loci, and this can favour the spread of reduced recombination around sex determining regions. Given that sex-linked regions are frequently repetitive and highly heterozygous, few complete Y chromosome assemblies are available to test these ideas. The guppy system (*Poecilia reticulata*) has long been invoked as an example of sex chromosome formation resulting from sexual conflict. Early genetics studies revealed that male colour patterning genes are mostly but not entirely Y-linked, and that X-linkage may be most common in low predation populations. More recent population genomic studies of guppies have reached varying conclusions about the size and placement of the Y-linked region. However, this previous work used a reference genome assembled from short-read sequences from a female guppy. Here, we present a new guppy reference genome assembly from a male, using long-read PacBio single-molecule real-time sequencing (SMRT) and chromosome contact information. Our new assembly sequences across repeat- and GC-rich regions and thus closes gaps and corrects mis-assemblies found in the short-read female-derived guppy genome. Using this improved reference genome, we then employed broad population sampling to detect sex differences across the genome. We identified two small regions that showed consistent male-specific signals. Moreover, our results help reconcile the contradictory conclusions put forth by past population genomic studies of the guppy sex chromosome. Our results are consistent with a small Y-specific region and rare recombination in male guppies.

## Introduction

Sexual conflict is thought to be one of the main drivers for the evolution of non-recombining sex chromosomes regions (Bachtrog 2013; Ponnikas et al. 2018). If a sexually antagonistic polymorphism (where at least one allele is beneficial in one sex and not the other) becomes established in a region linked to the sex determining locus (SDL), closer linkage between these two loci reduces conflict, allowing each sex to approach its different fitness optimum (Rice 1987). Since whole genome sequencing is now possible for a wide range of species we can begin to directly test these hypotheses by identifying the SDL and linked genomic regions. Additionally, by studying the sequences of young, only slightly differentiated, sex chromosomes, we can begin to distinguish causes and consequences of sex chromosome formation (Charlesworth 2019). Teleosts are particularly exciting in this regard because they have undergone frequent turnovers of their SDLs and often have young sex-linked regions (e.g. the medaka, *Oryzias latipes* and its congeners (Kondo et al. 2004)). However, identifying sex-specific regions of a genome with short-read sequencing is challenging, given that sex chromosomes and their sex-linked regions tend to accumulate repetitive sequences and are heterozygous for the SDL and potentially other fully sex-linked genes. Here, we present an improved genome assembly for the Trinidadian guppy (*Poecilia reticulata*) using long-range sequencing data and chromosome contact information in order to more fully explore the structure and evolution of their sex chromosomes. We then use the newly assembled sex chromosome and broad population sampling to detect sex differences across the genome.

The guppy has long been considered a species where sexual conflict may be important in sex chromosome evolution. This species has XY sex determination and is highly sexually dimorphic: males—the heterozygous sex (Winge and Winge 1927)—display complex colour patterns across their bodies and fins, while females are drab. Colour is used to attract mates but it also attracts the notice of predators. Guppies inhabiting rivers in the Northern Range of Trinidad are particularly well-suited for comparative study of colouration because colour patterning varies with predation regime. Here, the rivers are punctuated by waterfalls that define distinct predator assemblages: below waterfalls, guppies coexist with many predators (high predation = HP) while upstream they are relatively free of predation (low predation = LP). Guppies have evolved in response to their predators, displaying larger and more diverse colour spots in LP than in HP populations across multiple rivers (Endler 1995; Magurran 2005). Furthermore, the Northern range mountains of Trinidad act as barriers between rivers, and populations in different rivers are highly differentiated (Willing et al. 2010; Fraser et al. 2015). Therefore, the separate rivers represent a natural experiment, with guppies adapting to low predation regimes independently, providing the opportunity to study repeated evolution.

Early genetic work on the guppy revealed that male pattern formation genes are disproportionately Y-linked, with most colour traits co-segregating with the SDL or recombining only at very low frequencies (Haskins et al. 1961; Lindholm and Breden 2002). Even more intriguingly, Haskins and colleagues (1961) uncovered evidence for recombination between the sex chromosomes, and a difference in sex-linkage between predation regimes: the *Sb* colour factor was completely Y-linked in an Aripo river HP population, as testosterone-treated, sex-reversed females never displayed this trait, while sex-reversed females from an Aripo LP population often did. These observations demonstrated that the factor is present on the X chromosome, which suggests a history of crossing over between the Y and X in the LP population. This pattern was subsequently shown to apply across multiple populations from multiple rivers: LP females consistently displayed colouration following testosterone treatment more often than did HP females from the same river (Gordon et al. 2012). This result has been taken as evidence for a reduced rate of X-Y recombination in HP populations (Gordon et al. 2012) or, perhaps more simply, as evidence for weakened selection against colouration factors in LP sites (Charlesworth 2018).

Genome sequencing should help identify the SDL and help us understand its linkage with colour pattern genes in guppies. Independent genetic mapping studies have mapped the SDL to linkage group 12 (LG12) across multiple populations of guppies (Tripathi et al. 2009a; Bergero et al. 2019) and its sister species *P. wingei* (Lindholm and Breden 2002). Chromosome *in situ* hybridization studies revealed variation among populations in the size and heterochromatin content of a Y chromosome region towards the terminal end of LG12 (Nanda et al. 2014). Whole genome sequencing across populations of guppies, however, yielded unclear conclusions about the extent of the fully sex-linked region in Trinidadian guppies. Wright et al. (2017) detected sex-specific differences in SNP density and mapping coverage on LG12 and concluded that the non-recombining portion of the Y includes two evolutionary strata (fully sex-linked regions differing in ages) spanning 15-25 Mb on LG12 (Wright et al. 2017). These authors further concluded that the fully Y-linked region is larger in LP than HP populations, the opposite of the difference in recombination outlined above. However, another population genomics study found no evidence for such strata; instead the results suggested that the completely non-recombining part of the Y chromosome is small, perhaps a single gene (Bergero et al. 2019). Cytogenetic studies on crossovers in males (Lisachov et al. 2015) and linkage maps derived from multiple intrapopulation crosses (Bergero et al. 2019), observed male recombination events only near the ends of the chromosomes, in autosomes as well as the in the sex chromosome. Low recombination rates in male meiosis and non-recombining regions were also previously observed in genetic F_2_ crosses (Tripathi et al. 2009b). This pattern of sex-specific recombination would enhance linkage between the SDL and sexually antagonistic alleles even if they are physically distant from each other in both predation regimes (Bergero et al. 2019).

Both population genomic studies were limited by the available reference genome for guppies, which was assembled from short-read sequences from a female guppy (Künstner et al. 2016). Short-read sequencing technologies provide considerable challenges to assembling whole chromosomes and this can be particularly true for sex chromosomes, which are often densely populated with repeats. Repetitive regions can cause unresolved assembled regions of the genome, particularly if the repetitive region is longer than the library insert length (Alkan et al. 2011). Repeat elements, including transposable elements (TEs), are expected to be enriched on young sex chromosomes, and enrichment has been detected in *Drosophila miranda* (Bachtrog 2003) and in teleosts (Chalopin et al. 2015). Moreover, TEs have been implicated in the movement of SDLs from one chromosomes to another (e.g. salmon (Faber-Hammond et al. 2015; Lubieniecki et al. 2015), and in the regulation of nearby genes, including the sex-determining genes themselves (e.g. medaka (Herpin et al. 2010), reviewed in (Dechaud et al. 2019)). Long-range sequencing has the promise to sequence across repetitive regions, but even these remain limited by maximum read lengths. Chromosome contact information (e.g. Hi-C) identifies long-range chromosome contact information without prior marker selection and can therefore improve contiguity of assemblies by placing (or scaffolding) contigs along chromosomes (e.g. (Lieberman-Aiden et al. 2009).

Here, we report a new guppy reference genome assembly that uses a male sample, and that employs both long-read PacBio sequencing and HiC-based chromosome contact information. We show that our new reference assembly closes gaps and corrects mis-assemblies in the short-read female-derived guppy genome sequence. Importantly, for LG12, we have captured GC- and repeat-rich regions not previously assembled. Using this improved reference genome, we then employed a population genomic approach to detect differences between males and females across the sex chromosome. We identified two small regions on LG12 that show consistent evidence for sex-linkage. These advances demonstrate the importance of drawing on new techniques for improving existing genomic resources and highlight the benefits of combining high-quality assemblies with population genomic approaches. Furthermore, our results help reconcile the contradictory conclusions of past studies of guppy sex chromosome evolution that relied on a female reference genome, and suggest new directions for evolutionary genetic study of sex chromosome evolution in guppies.

## Results

### A long-read assembly of a male guppy genome

We assembled the genome of a single male from the same inbred Guanapo strain that was used for the original female reference genome (Künstner et al. 2016). Our approach integrated 50 Gb of PacBio long read data and 40 Gb of Hi-C chromosomal contact data into a phased, diploid assembly with a total size of 745 Mb (Table S1: analysis was done on the slightly longer phase0 if not otherwise stated, see below for a description of the two phases). First, complete chromosomes were reconstructed using three previously generated genetic linkage maps ((Tripathi et al. 2009a): GM1; (Künstner et al. 2016): GM2; (Bergero et al. 2019): GM3, for LG12 only). Assembly misjoins in 11 contigs were corrected by splitting at breakpoints inferred in genetic maps GM1 and GM2 and inter-chromosomal contact domains detected from remapped Hi-C data. The final chromosome set totals 698 Mb and integrates 93% of all assembled contigs (Figure S1; Figure S2, Figure S3, Figure S4). BUSCO completeness was 94.9% (93.7% single-copy, 1.2% duplicated), with 1.5% fragmented and 3.6% missing, based on a total of 3640 BUSCO groups.

This new male genome assembly (named MG) is both longer and more contiguous than the previously released female guppy genome, FG (total scaffold length 745 Mb in MG vs total contig length 663 Mb in FG and total scaffold length of 731 Mb in FG) and has a much higher N50 contig length of 7,936,040 bp versus 35,577 bp for the FG. Comparing the mapping statistics of short reads from natural guppy populations to both genome assemblies, we found an increase in the mean number of paired reads that mapped properly (95.4% vs. 91.4%, Table S2). The increase was similar with reads from either sex and higher for reads from individuals collected in the source river, Guanapo, than from individuals in the two other rivers sampled.

Alignment of RNA-seq data to the softmasked genome (28.3% softmasked) yielded alignment rates between 86.0% and 93.8%. The BRAKER2 annotation pipeline used an initial set of 83,491 open reading frames without internal or terminal stops. An initial dataset, from *Xiphophorus maculatus*, (a closely related species) used for calculating the overlap-hit ratio (OHR: see methods), contained 36,613 proteins, which collapsed down to 27,791 at a ≥90% identity threshold after CD-HIT clustering. Orthology comparison between the *P. reticulata* and *X. maculatus* proteins using BLAST with an e-value threshold of 1e-5 and an OHR ≥95% yielded a final annotation set of 18,227 proteins, and subsequent functional annotation using EggNOG using an e-value threshold of 1e-10 resulted in a final set of 15,263 protein coding genes. Repeats were annotated with RepeatMasker (Table S3). The overall GC level was 39.7%, and 9.9% of the genome (73,979,443 bp) was masked for repeat content (Figure S5).

### Whole genome alignment

The new male genome assembly was mostly syntenic with the previously published female genome assembly, and also with the genome of the closely related platyfish, *Xiphophorus maculatus* (Figure 1, Figure S6, Figure S7). However, many chromosomes are arranged differently in the new versus the previous assembly, including LG12, where the new assembly is inverted across the region from 11 Mb to 20 Mb (Figure 1). As the new orientation is supported by multiple genetic maps and largely corresponds to the platyfish assembly, this region was probably mis-assembled in the previous guppy genome sequence. Similar improvements of synteny with the platyfish assembly are found across the genome (Figure S7).

**Figure 1:**
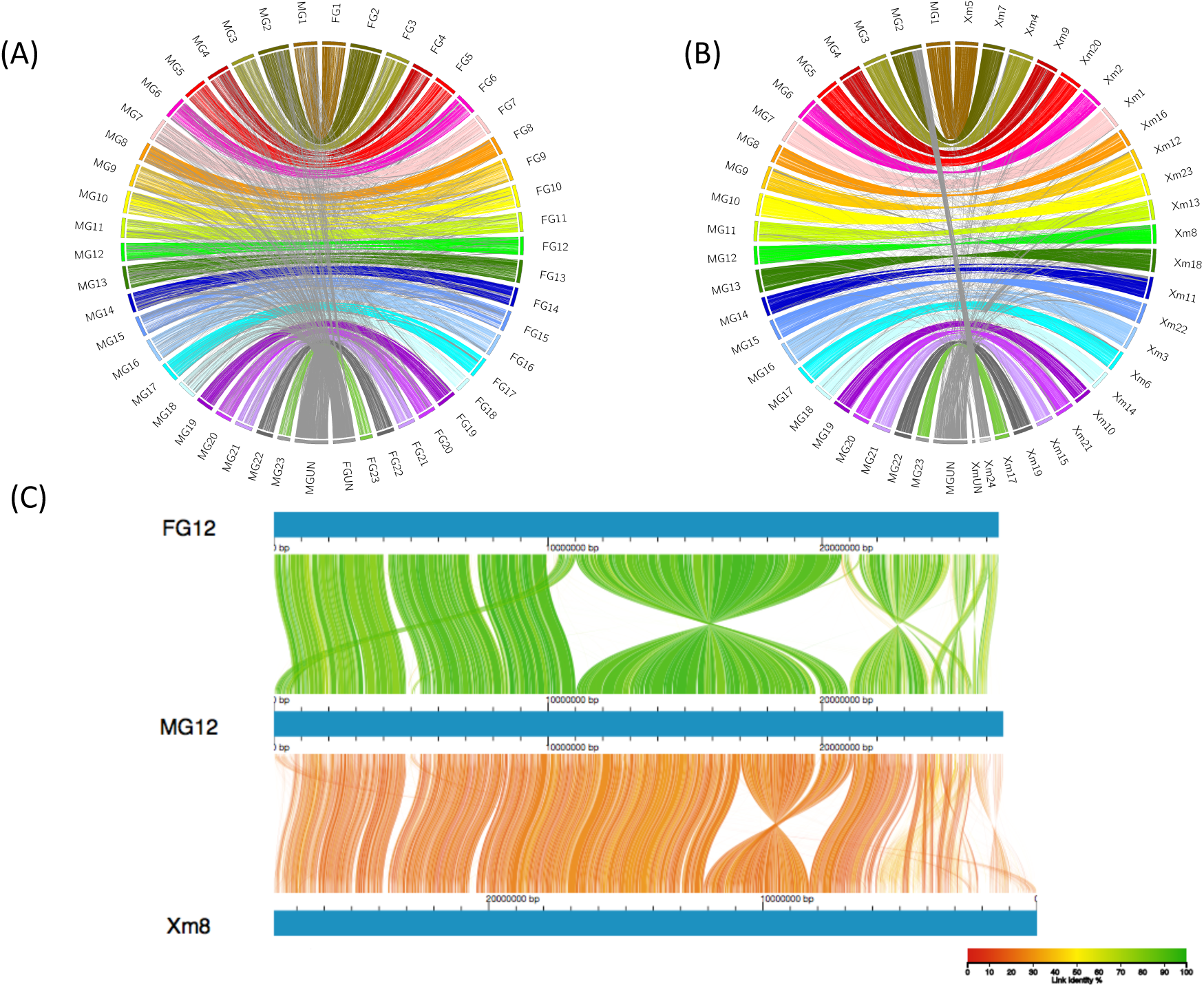
Whole-genome alignments of assemblies. (A) Whole-genome alignment between new male guppy genome assembly (MG) and a previous Illumina-based female guppy genome assembly (FG). (B) Whole-genome alignment between new male guppy genome (MG) assembly and *Xiphophorus maculatus* genome assembly (Xm). Links are coloured by chromosome, with between-chromosome and unplaced scaffold links coloured grey. (C) Genome alignment of LG12 between the two guppy assemblies and chromosome 8 on *Xiphophorus maculatus*. Colour denotes percent similarity of links (see legend).

### Examination of LG12

The motivation for a new male-based assembly was to investigate sex chromosome evolution. We therefore closely examined LG12’s gene content. In total, 14 contigs were placed onto LG12 (Figure 2). Placement was based on 29 SNP markers, 283 RAD-seq markers, and an additional 12 microsatellite markers from (Bergero et al. 2019) (Figure 2), as well as Hi-C information. A few contigs showed evidence of being misplaced in the release genome assembly according to the Hi-C data and therefore were manually re-ordered (see supplemental methods for description, Figure S8). Also, of note, Contig IV (at approximately 5 Mb) has only 2 SNP markers, one which places the contig at 5Mb and one marker that places it at the distal end of LG12, yet, contig IV has high sequence similarity to contig XII (see below), implying that it could be assigned equally well to either region and therefore not informative for placement. We next examined LG12 for gene and repeat features (Figure 2). Similar to most other chromosomes (Figure S5), LG12 showed an increase in GC content at the assembly end, most notably at 24.5 - 25.5 Mb (contig XIII) (Figure 2b). Particularly high repeat density was also apparent around 5 Mb (contig IV) (Figure 2c).

**Figure 2:**
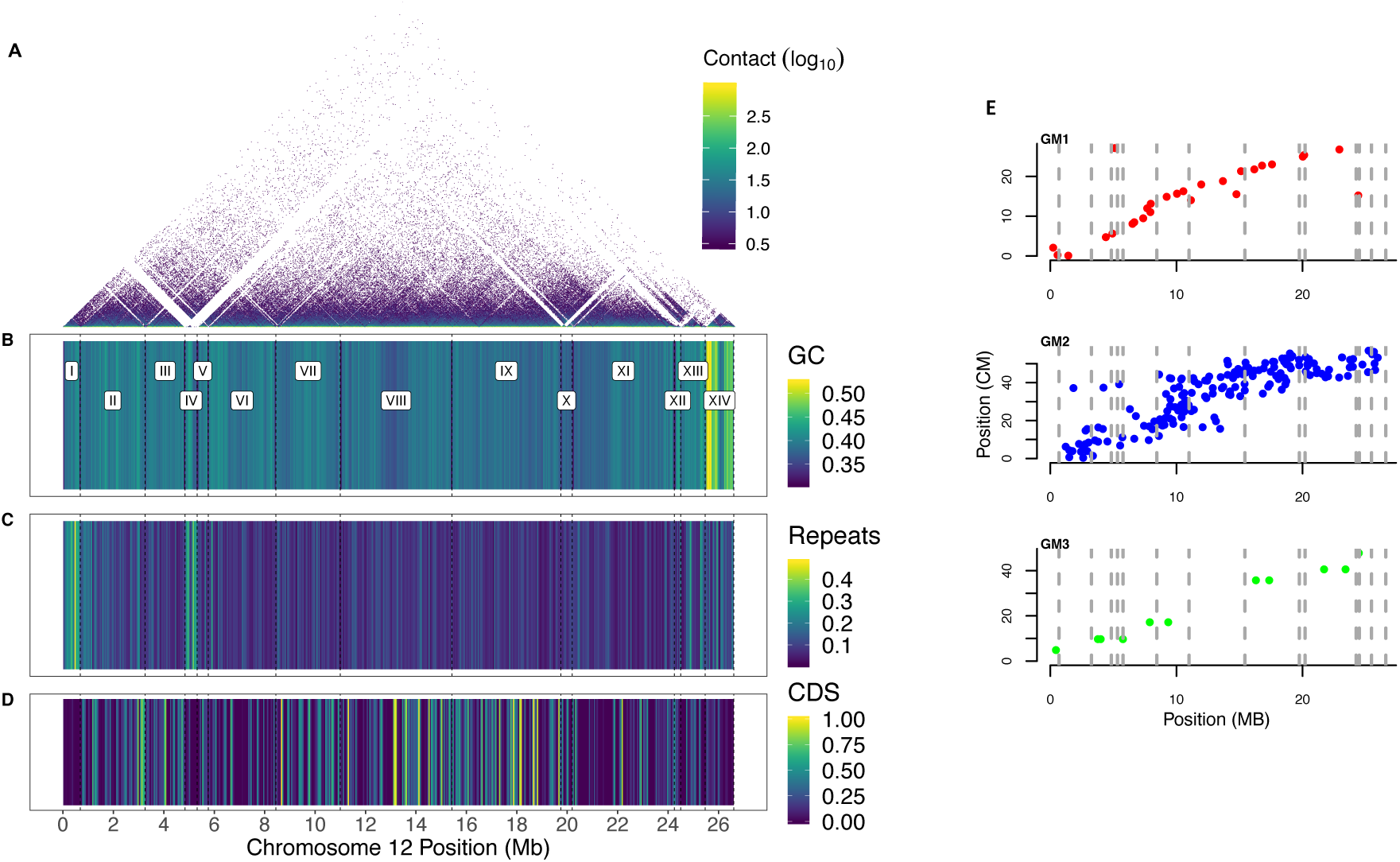
Assembly of LG12 from the male genome, with each contig being delineated with dashed lines. (A) Hi-C contact, (B) percentage GC content, (C) percentage repeat content, (D) percentage of CDS (B-D in 50 kB windows), (E) Genetic linkage maps along LG12, with each marker’s position shown on the MG genome in centimorgans (cM) for GM1, GM2, and GM3.

The genome assembly was phased using Falcon unzip, although each phased contig is arbitrarily placed into one chromosome scaffold or the other, so that the assigned phases do not necessarily correspond with the actual haplotypes across the entire chromosome in the sequenced individual. The phases should not be interpreted as X-Y haplotypes; the phases are a data structure without biological meaning beyond the contigs on each phase being mutually exclusive. We can, however, examine differences between each pair of phased contigs (Table S4). A first observation is that most contigs are of similar size in both members of the pair, with only Contig IV differing in size by >100kb (its phase0 contig includes a region named ‘C2’ not present in phase1, see below and Figure 4A for a full description of this region). Second, SNP variation and SVs are evenly distributed across LG12 (Figure S9). Notably, a large inversion exists between the phases (size = 973,876bp, beginning at 25.84Mb).

**Figure 3:**
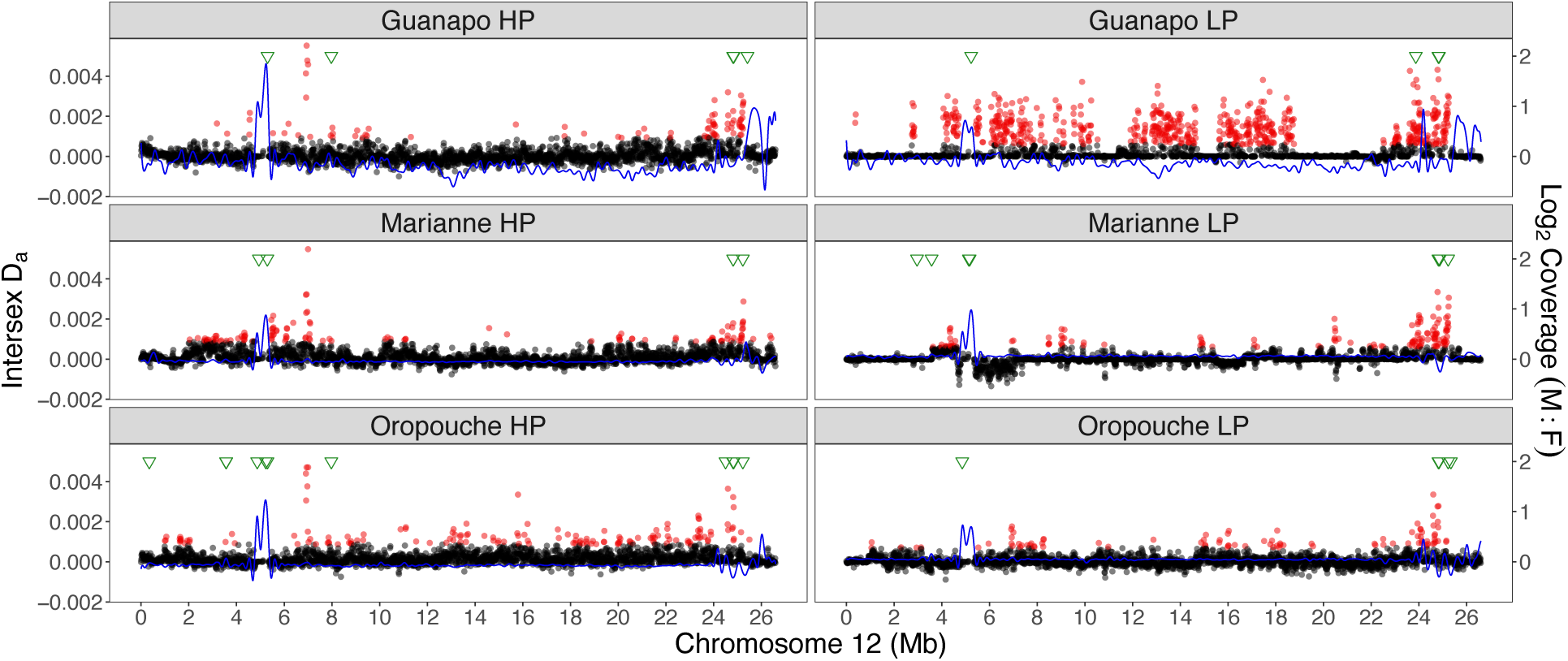
Population genomic analysis for sex-specificity of variants on LG12 in six natural populations. Each population is shown in a panel, with high-predation (HP) populations on the left and low-predation (LP) populations on the right. D_a_ values are shown as black dots, with outliers in red. Coverage differences between the sexes are fitted with a spline, shown in blue. Finally, the positions of male contigs assembled from male-specific assembled k-mers (y-mers) are indicated along the top in green triangles.

**Figure 4:**
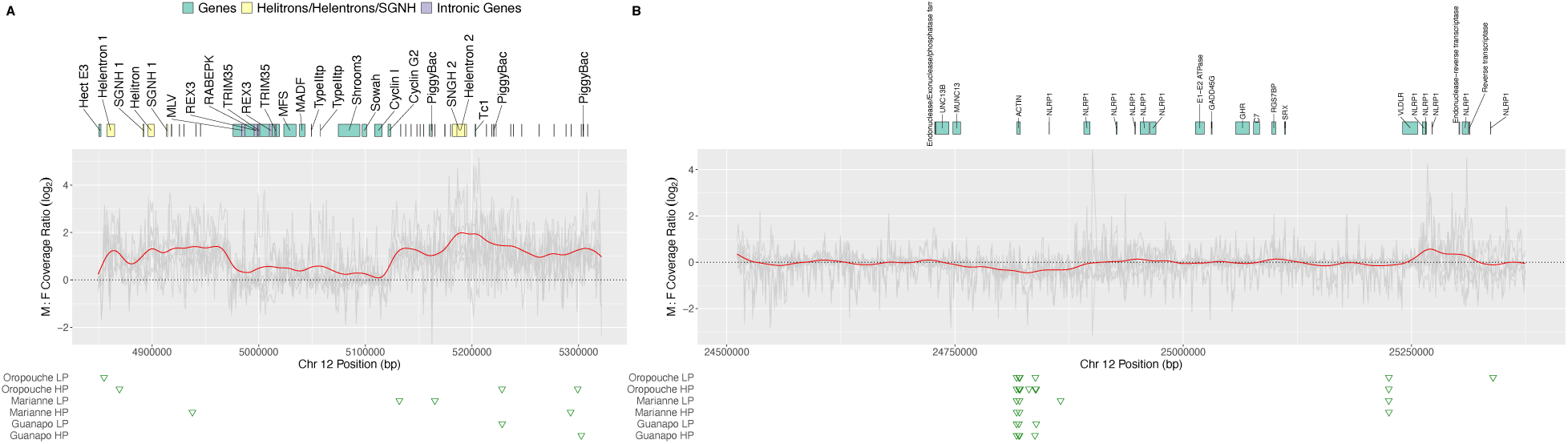
Detailed investigation of candidate regions on LG12 that are fully sex-linked. (A) Region 4.8-5.2 Mb, (B) region 24.5-25.4 Mb. Grey represents each population’s male/female coverage ratio, red the mean coverage ratio. Gene annotations are shown above as blocks above, male specific contigs for each population are shown in green triangles below.

### Population genomics genome scans

To identify the male-specific region of the Y chromosome, we took a population genomics approach, sampling males and females from six populations representing all three major drainages in Northern Trinidad (Table S5, Figure S10). We tested each population with three approaches that can detect sex differences: comparing read coverage and SNP diversity (F_ST_ and D_a_ = net D_XY_ (Nei 1987)) between the sexes, and searching for Y-specific k-mers (‘y-mers’).

We first scanned the genome for consistent differences in coverage between the sexes in 10 kb windows, using our short reads from individuals from natural populations: this can detect sex-linked regions in which males are hemizygous, or sequences of Y- and X-linked regions that are highly diverged. Note that we expect Guanapo populations to be best suited for the coverage test as the reference genome sample was derived from a Guanapo site. Consistent with previous results (Bergero et al. 2019), coverage was generally uniform across the genome, and mean male/female coverage difference was not consistently higher on LG12 compared to other chromosomes (Figure S11). Female-biased coverage was variable across populations (numbers of female-biased 10 kb windows in different populations samples were as follows: GH=1,109, GL=240, MH=14, ML=24, OH=59,OL=43). No window showed consistent female bias in all six populations, but one window was shared by four populations, and two were shared among three; however, none of these was on LG12 (Table S6).

There were considerably more male-biased windows across the genome (total numbers of male-biased 10 kb windows: GH=2,306, GL=1,230, MH=377, ML=269, OH=626, OL=517), and there was also more overlap among the populations: 57 windows were found in all six populations, 151 in five populations, and 154 in four populations (Table S6). Of the windows shared by all six populations, most (45 windows) were on unplaced scaffolds and only three were on LG12, clustered on contig IV (between 4.9-5.3 Mb, Figure 3). An additional 11 10kb windows were detected within this region that exhibited male-biased coverage in any 5 populations. The remainder were on LG18 (n = 6) and LG5 (n = 3). Overall, coverage patterns were more similar within river drainage than the predation regime.

To look for sex differences in SNP frequencies we first estimated F_ST_ between the sexes in 10kb windows along the genome. A value of 0.3 is expected for a fully sex-linked XY SNP site or region (heterozygous in all males and homozygous in females). Genome-wide F_ST_ between the sexes was low within all populations (GH = 0.022, GL = 0.014, MH = 0.019, ML = 0.017, OH = 0.020, OL = 0.017). No chromosomes showed elevated F_ST_, except LG12 in GL (F_ST_ = 0.11, see Figure S12). Elevated F_ST_ was evident across the whole chromosome in the GL population and was due to greater nucleotide diversity in males than females, as expected for sites or regions in linkage disequilibrium with the SDL (Figure S13). Values of F_ST_ above 0.3 varied across the six populations, and were highest in GL and low in OH and OL (the numbers of windows with F_ST_ > 0.3 were: GH = 174, GL = 661, MH = 135, ML = 189, OH = 23, OL = 38). Windows with elevated F_ST_ values were shared between at most three populations, and involved one window on LG12 and three windows on unplaced scaffolds (Table S7). We also scanned LG12 for F_IS_ outliers. While, F_ST_ and F_IS_ are closely related, F_IS_ has the advantage that it can detect outliers that are due to males being more heterozygotic than expected based on the variant frequencies, which in turn supports sex linkage (complete or partial). However, like our F_ST_ results, there were no consistent F_IS_ outlier windows across LG12 (Figure S14).

We also estimated net sequence divergence between the sexes in the same 10 kb windows. F_ST_ between the sexes depends strongly on diversity within the two sexes (Charlesworth 1998), we therefore used a net divergence measure, corrected by female diversity, D_a_, which can detect regions with higher diversity in males than females. Importantly, this measure is sensitive to regions enriched for low-frequency, male-specific variants, due for example to diverse Y-linked haplotypes segregating within populations (Bergero et al. 2019), which would produce low F_ST_ values. For each population, we identified candidate windows with D_a_ values greater than the genome-wide mean plus 3 standard deviations (akin to a 99% confidence interval). LG12 had the highest average D_a_ in five populations, and had the second highest average in the ML population (Figure S15). Six windows had outlier D_a_ windows in all six populations: four were located at the distal end of LG12 (between 24.5 and 24.9 Mb) and the other two were found between 6.94 and 7.01 Mb of LG12 (Table S8, Figure 3).

Given the difficulty of detecting SNP differences when mapping is low in some samples, we also applied a k-mer assembly approach. First, we identified all 12 bp k-mers that could be found in all males, but were missing in all females in all six populations. The reads containing such y-specific k-mers or ‘y-mers’ were then reassembled for individual populations to assemble ‘male-specific contigs’, and finally realigned to our genome assembly. In total we assembled a mean of 132 male-specific contigs per population, with a mean total length of 64,528 bp (Table S9). These contigs were distributed unevenly across our reference genome assembly (Figure S16, Table S9) but LG12 was consistently over-represented, with >5% of total contig length recruited to LG12 in all populations (Table S9). These contigs are mostly clustered in two regions, at 5 and 24-25 Mb (Figure 3). Autosomal regions also recruited male contigs, but less consistently across populations (Figure S16). Overall, we identified only ten 100 kb windows where all six populations had male-specific contigs (LG12 24-25 Mb, LG18 27-28 Mb, LG5 32-33 Mb and six unplaced scaffolds [000173F, 000175F, 000189F, 000195F, 000246F, 000275F, and 000281F], Figure S16). Not surprisingly, most of these windows were also male-biased candidates based on coverage (Table S6). The male-biased coverage regions on LG12 (4-6Mb) also recruited contigs in all populations, but not in directly overlapping windows (Figure 3).

Taken together, two regions appear most likely to be candidates for full sex-linkage: contig IV and contig XIII of LG12. Both regions contained y-mer contigs in all six populations, and contig IV had consistently high male-biased coverage, while contig XIII of LG12 had windows with consistently high D_a_ values. The same criteria (intersection of male biased coverage and recruitment of male contigs or consistent D_a_ outlier) also pointed to four unplaced scaffolds (000173F, 000175F, 000189F, 000246F) and one autosomal region (LG5 32-33 Mb) as apparently partially sex-linked; gene annotations for these regions are provided in Table S10.

### Further investigation of candidate regions

#### Contig IV (LG12 4.8-5.2 Mb)

Closer inspection prompted us to divide this contig into three distinct regions; a repeat-rich region (C1), a gene island (G1), and another repeat-rich region (C2) (Figure 4A). C1 and C2 show evidence for higher coverage in males than females and presence of male-specific contigs, suggesting complete sex-linkage. As expected for regions with variable mapping rates due to the male-biased coverage, C1 and C2 lack SNPs after filtering, and F_ST_ or D_a_ could not be estimated here.

Given its repetitive content, we annotated this contig manually. First, using the reference transcriptome from Sharma and colleagues (2014), we retrieved a preliminary set of 273 blastn matches that we then reduced to 71 non-redundant matches via an all-against-all blastn search, retaining only the longest transcript within each cluster of hits. Querying these transcripts in Ensembl resulted in annotations for 30 transcripts. Sequences with matches tended to be considerably longer (median = 975 bp; interquartile range = 424-1,532 bp) than those without (median = 216; interquartile range = 183-297 bp). Nineteen of the 30 annotated transcripts matched known repetitive elements (transposons, retrotransposons, helitrons, or retroviruses); these mostly mapped to the ends of contig IV. The remaining 11 were considered putatively “non-repetitive” protein-coding transcripts (Figure 4A). One transcript, which encoded a predicted hAT transposon C-terminal dimerization domain, mapped to 27 unique positions throughout the highly repetitive regions; separate analyses showed that XIR and XIR-like repetitive elements also repeatedly mapped to these stretches. XIR repeats are particularly interesting because they have been shown to be Y specific in platyfish (Nanda et al. 2000; Volff and Schartl 2001).

The gene annotation of contig IV revealed similarity with Contig XII, which is assembled near the 24 Mb point of LG12 (Figure 2). A NUCmer alignment (with flags “--maxmatch -l 75”) detected a region of similarity of roughly 0.15 Mb between the two contigs, encompassing the gene-rich region (G1) of contig IV (Figure S17). A few repetitive elements also mapped within G1, most notably a MuLV retrovirus and two copies of a REX3 nonLTR retrotransposon; both mapped to a region near the upstream boundary of the duplicated stretch where synteny was clearly disrupted between contigs IV and XII (Figure S17). The heavily disrupted region between the contigs that contains the MuLV and REX3 sequences corresponds to a large intron of a RABEPK-like gene, within which we also found two copies of a short TRIM35-like gene (Figure S18). The corresponding intronic region appears to be shorter in contig XII, lacking MuLV and both copies of REX3, and having only a single TRIM35-like gene (with two more copies roughly 0.1 Mb upstream of the start of Contig XII). The RABEPK-like gene is followed by genes for an MFS transporter, a MADF-domain containing protein (a putative transcription factor), Shroom3, a Sosondowah ankyrin repeat domain family (SOWAH)-like protein, Cyclin I (which is partially triplicated in Contig XII), and Cyclin G2.

Many of these genes have previously been noted as being present on the guppy and platyfish sex chromosomes. Tripathi and colleagues (2009b) found a sex-linked SNP within a Cyclin G2 gene. Cyclin G2 was also found to have male-biased expression by Sharma et al. (2014). The most terminal marker in GM1 (marker 229) was used to isolate BAC 36_H23 (Tripathi et al. 2009b). This BAC labeled the distal ends of the X and Y chromosomes in several guppy populations (Nanda et al. 2014). Blasting this BAC to the MG sequence shows that it maps to an unplaced scaffold (00250F) and to the first cyclinI gene on contig XII, potentially placing this contig on LG12. Notably, this unplaced scaffold also has D_a_ outliers in 5 of 6 populations (Table S8). The potential role of cyclins in multiplication of germ cell precursors might be worth investigation (Schulz et al. 2010). Morris and colleagues (2018) assembled a male-specific k-mer contig that also contained a TRIM39-like gene. TRIM35 and TRIM39 have the same domain architecture (C-IV-1, (Hatakeyama 2017). However, our reference assembly includes many copies of TRIM genes (we found 126 homologs distributed across the genome based on a blastn search). Both the TRIM and cyclin genes are interesting because it is at these genes where the duplicated regions differ (Figure S17). Using a PCR presence/absence test the TRIM35-like gene was found to be associated with sex in only some of our populations (Table S11).

It is notable that only the repetitive regions C1 and C2 show Y-like male:female coverage differences, but not the gene-rich region G1 (Figure 4A). These repetitive regions were also very similar to other regions across the genome, particularly to unplaced scaffolds. When unplaced scaffolds were aligned individually to all chromosome assemblies using minimap2, and after alignment blocks shorter than 10,000 bp were filtered out, LG12 was left with more alignments (N = 151) than any of the other 22 chromosomes (4 ≤ N ≤ 90). Most of these LG12 alignments (107/151) were clustered on contig IV, aligning to 51 unique scaffolds. Visualisation of the alignments highlighted two contig IV regions with overlapping high-quality alignments (above the median Smith-Waterman alignment score for contig IV alignments of 21,584). These clusters aligned to a 25 kb region (LG12:4,942,488-4,967,664 bp [C1]) and a 48 kb region about 200 kb away (LG12:5,175,028-5,213,274 bp [C2]) (Figure 4A). Searching the full genome with these sequences identified additional autosomal sequences on LG18 (C1 + C2), LG15 (C1), and LG2 (C1) with similarity to these two LG12 regions. The scaffolds that make up these clusters were extracted and re-aligned to contig IV. Inspection of the alignments for structural variation using MUMmer 4 (Kurtz et al. 2004) (Figure S19) detected minimal variation, except for an inversion in the C2 related sequence on LG18 (27,568,621-27,588,735 bp). Many of these aligned regions also exhibited Y-like male:female coverage ratios and account for at least one of our ‘other candidates’ (000173F) and to a coverage candidate (LG18 27-28 Mb, Table S6).

Both the C1 and C2 sequences were annotated as including helitron transposons and a SGNH hydrolase domain-containing gene (Figure 4A). Further examination of these genes by searching the NR database revealed similarity to the texim/SGNH genes from platyfish (Tomaszkiewicz et al. 2014). The relevant regions within C1 best matched type 2/3/4 texim, whereas C2 best matched type 1/Y texim genes (Tomaszkiewicz et al. 2014). The platyfish teximY and texim1 sequences, which are both approximately 1300 bp long, differ by only a few bp (Tomaszkiewicz et al. 2014). The platyfish Y chromosome (Xm LG21, corresponding the guppy LG20) has a higher helitron density than the X chromosome (Zhou et al. 2006). Y-chromosome located texim/SGNH hydrolase genes involved in spermatogenesis have apparently been captured by helitrons (teximY) (Tomaszkiewicz et al. 2014). Using qPCR, we found that the copy number of these texim-type/SGNH hydrolase genes was higher in males than females in five of the six populations sampled (Table S12). We were unable to find any teximY-specific sequences unique to contig IV that could be used as a sex-linked marker.

The repeats in the C1 and C2 regions do not have any good matches in the previous female guppy genome assembly (Künstner et al. 2016). The closest partial matches were observed on unplaced scaffolds 853 (with C1) and 221 (with C2). The latter included a good blast hit (E=0) for the texim1 region of C2.

### Contig XIII (24.5 - 25.4 Mb)

Although the SDL has been mapped to the distal end of LG12 previously (Tripathi et al. 2009a), the evidence for Y-linkage was less clear, since we found no consistent read coverage or F_ST_ signal differentiating the sexes. However, all populations show male-specific y-mer contigs that map to around 24.82 Mb, regions with elevated D_a_ from 24.59 - 24.61 Mb, 24.78-24.79 Mb, and 24.82-24.83 Mb, and a marginal dip in male coverage at 24.8 (Figure 4b). Taken together, these results suggest a recent increase in differentiation between the X and Y in this region. In terms of gene content, this region consists of an actin gene followed by multiple copies of NLRP1-related genes. NLRP1 is part of the inflammasome, an important element of vertebrate immunity (Mitchell et al. 2019). NLRP1-related genes were also found on two of our candidate unplaced scaffolds (000173F and 000189F, Table S10).

Multiple lines of evidence from previous work had identified this region as the one most likely to include the guppy SDL. Tripathi and colleagues (Tripathi et al. 2009a) located the SDL distal to the marker 229, which is at 24.4 Mb in this new assembly. We find few and inconsistent F_IS_ or F_ST_ signals in this region in our results (Figure S14), and others reported similar findings for a sample from a captive population (Bergero et al. 2019). Dor and colleagues (2019) proposed that the SDL is within the distal 5.4 Mb of LG12, based on an association between sequence variants and sex in ornamental guppies. This region is rearranged in the male assembly compared to the previous female assembly (Figure S20). Of particular interest are the candidate genes proposed by Dor and colleagues (2019); stoml2 (at 24,680,972-24,686,396 bp), which was not annotated by our pipeline but found through an Ensembl comparison, and gadd45g (25,035,744 - 25,039,913 bp) (Figure 4B). Neither gene shows a consistent signal of sex differentiation, but the region between the two genes includes our candidate region around 24.82 Mb. Therefore, these candidate genes are probably within a region associated with sex in the ornamental population studied by (Dor et al. 2019), but not in wild populations.

## Discussion

The Trinidadian guppy, *Poecilia reticulata*, is a promising system for study of the early stages of sex chromosome evolution, but this work has been hampered by the lack of a highly contiguous genome that includes the male-specific Y chromosome. We present a new chromosome-level reference genome assembly for the Trinidadian guppy, *Poecilia reticulata*. PacBio single-molecule real-time sequencing reads and Hi-C chromosome contact data supported the assembly of repetitive and GC-regions that were not previously resolved. After correcting a number of apparent mis-assemblies, the assembled chromosomes are now substantially more syntenic with those of the related platyfish, *X. maculatus* (Schartl et al. 2013). This new assembly has been particularly useful for examining the sex chromosome (LG12), and we highlight two small regions that are consistently associated with sex in samples of paired HP and LP natural populations from three river basins. Within the river basins, we find no evidence that HP and LP populations differ in the size of the sex-linked region, contrary to predictions and past observations. Neither do we find evidence of extensive evolutionary strata of complete sex linkage. We conclude, in agreement with Bergero and colleagues (2019), that any non-recombining fully Y-linked region must be small. Our detailed analysis provides new SDL candidates for this region, which were missed in previous analyses that relied upon the female reference genome.

Our most promising region—situated at approximately 5 Mb on LG12— contains candidate loci that have been previously mapped to the Y chromosome in guppy and platyfish. The region consists of two highly repetitive stretches that surround a genic region that itself is homologous to a region at 24 Mb on LG12. The two repeat-rich regions show similarity to many unplaced scaffolds, and to regions within LG18 and LG4. Our candidate region is not present in the previous female genome assembly, and thus has not been detected in genome scans before. Reads originating from this region in previous genome scans were probably unmapped or even mis-mapped elsewhere in the female assembly, either to the duplicated genic region on LG12 near 24 Mb or to related sequences elsewhere in the genome. The most distal repeat-rich region includes our strongest evidence for increased male coverage; it is not present in the alternate phased assembly, and contains teximY genes, which have previously been found on the platyfish sex chromosome (Tomaszkiewicz et al. 2014). Notably, the platyfish sex-linked region is located on Xma21 (Schartl et al. 2013); it corresponds to guppy LG20, where we find no evidence of sex-bias.

Locating the SDL on a contig at 5 Mb of LG12 is contrary to genetic and cytogenetic studies that suggested it to be near the distal end of LG12. Our evidence for placing this contig at 5 Mb is admittedly mixed. The Hi-C signal is low in this region, likely because of low mappability due to the high number of repeats (Lajoie et al. 2015). Only two genetic markers were mapped here and one shows linkage to the other end of the chromosome (at 27.17 cM, Figure 2). Longer sequencing reads, or BAC sequences, could further resolve the placement of this region further, as well as that of the many unmapped scaffolds that show sequence similarity to contig IV.

Our second candidate region is at the distal end of LG12, where the SDL has been mapped previously. Here, our evidence for sex-linkage is higher net divergence between the sexes than elsewhere on LG12, and recruitment of contigs that were assembled from y-mers, male-specifc k-mers. However, we detected no consistent coverage differences between the sexes and no high F_ST_ between sexes in this region. Together, our results support the previous conclusion (Bergero et al. 2019) that divergence between the X- and Y-linked region is small, and that no obvious evidence for genetic degeneration along the Y chromosomes. An alternative possibility is that the sex-determining region includes additional repetitive sequences and highly divergent regions that were not assembled here. While there is no obvious nearby candidate sex-determining gene in our candidate region (∼ 24.82Mb, contig XIII), this region shows some overlap with the region identified by Dor and colleagues (2019), (Figure S20). The candidate contig is followed by a GC-rich region, corresponding to the end of the chromosome where recombination is known to be high and shows no evidence for sex-linkage (Bergero et al. 2019, Charlesworth et al. 2020).

Given that the SDL has been mapped to the terminal part of LG12, the absence of clear signs of male-specific variants (i.e., high F_ST_ or biased male:female coverage ratio) in this region is surprising. Y-linked regions often show low diversity (Cotter et al. 2016), which would lead to high male-female F_ST_ values. One possibility is that the region includes many different Y haplotypes, which would lower male-female F_ST_ values. By using a net divergence measure (D_a_) we detected signals of high male diversity relative to female diversity, resulting from many low-frequency male specific variants. Balancing selection for Y-linked colour variants could be responsible for maintaining many Y haplotypes, with evidence for negative-frequency dependent selection for guppy colour patterns coming from manipulating colour pattern frequencies in natural populations. In such experiments, Olendorf and colleagues (2006) found that males with a rare colour pattern had a survival advantage compared to common males. Similarly, Hughes and colleagues (2013) found that rare males had reproductive advantage. The hypothesis of multiple co-existing Y haplotypes is testable using long-read sequencing that can determine the phase of variants in the region, or phasing using sequencing of either mother-father-offspring trios or of individual sperm.

We next examine the controversy over the size and evolution of the Y chromosome in guppies. Wright and colleagues (2017) presented evidence for two evolutionary strata spanning the distal 10 Mb of LG12, while Bergero and colleagues (2019) did not find a clear region with sex-specific signals. Our new assembly updates these two strata substantially and can also explain inconsistent signals. First, the putatively younger strata (15-22 Mb in the previous female assembly) identified by Wright and colleagues (2017) was based on higher SNP density in males than females, although Bergero and colleagues (2019) found no consistently elevated F_ST_ in the population they examined. Our new assembly rearranges the region containing the putative stratum into non-consecutive regions (Figure 1C). We too find no consistent signal of high male diversity across these regions, nor do we find these regions differ between HP and LP populations. The putatively older stratum (22-25 Mb in the female assembly) was identified through coverage differences between the sexes by Wright colleagues (2017) but not by Bergero and colleagues (2019). In our new assembly, these sequences remain largely in place, albeit with some rearrangements (Figure S20). Our candidate region (∼24.8 Mb), is considerably smaller than that reported by Wright and colleagues (2017). The differences among these studies most likely stem from our increased sequence length and different bioinformatic filtering pipelines, as well as from the fact that read mapping to our male reference genome provides a more direct view of Y chromosome evolution than analyses based on the X chromosome. For example, our candidate region at 5 Mb shows homology with contig XII at 24 Mb, which corresponds to roughly 23 Mb in the female assembly. It is therefore possible that small differences in mapping filters could account for the differences in signals reported among studies.

Our approach of using a multiple population genomics scan was critical for excluding regions from further consideration, and for narrowing down candidate regions. Had we analyzed only one population, we would have come to very different conclusions. For example, the GL population showed differences between the sexes across most of LG12, which suggests that this up-river population was founded by a small number of females. Such a bottleneck would have reduced diversity, which can explain this population’s high F_ST_ values between the sexes across LG12, as F_ST_ estimates the times back to a common ancestor, relative to that within the sub-populations being compared (here, the two sexes). A bottleneck is consistent with females being able to store sperm from multiple males (Kobayashi and Iwamatsu 2002; Kurtz et al. 2004), and thus of a few or even just one female potentially founding a new populations.

Our genome scan analysis also detected sex-biased regions on autosomes, which could potentially indicate sexual antagonistic loci (e.g. a region on LG5 using a similar intersection of measures used to highlight our candidate regions). However, caution is needed, as individual-based simulations (Kasimatis et al. 2019) suggest that spurious signals of high F_ST_ between the sexes can result from sampling variance and low sample sizes. These simulations confirmed that antagonistic selection must be remarkably strong (e.g., be associated with high sex-specific mortality or fertility differences) to maintain differences between the sexes at autosomal loci, which seems unlikely for most species (Fry 2010).

Autosomal and pseudoautosomal regions with sex-biases in coverage can also be due to mis-mapping of reads. Bisseger and colleagues (2020) found that recent duplications and translocation onto the Y chromosome in stickleback are often responsible for incorrect inferences of candidate sexually antagonistic loci on autosomes when using genome reference assemblies that lack Y chromosome sequences. Indeed, multiply mapping reads could be at least partially responsible for the sex-biased signals that we found here, for example, a coverage outlier region found on LG18, and for the many unplaced scaffolds with high sequence similarity with our candidate region at 5 Mb on LG12.

Why are the two sex-biased candidate regions on LG12 not physically close in the male genome assembly? As previously discussed, the apparent distance between these two regions could be artefactual and disappear with additional sequencing and linkage information. That said, this arrangement is biologically plausible given the guppy’s recombination landscape. Specifically, Bergero and colleagues (2019) argued that associations between specific DNA sequences and sex could be maintained in separate regions of LG12 due to recombination in males being largely, if not entirely, restricted to the chromosome’s distal tips. Consequently, sexually antagonistic loci need not be physically linked to the SDL on LG12. We conclude that the Y specific region on LG12 is small, likely to be within one of our candidate regions. Additionally, our findings suggest that the SDL may not be physically close to sexually antagonistic loci situated on the Y chromosome, but may instead be genetically closely linked due to the rarity of crossovers in male meiosis (Bergero et al. 2019).

## Methods

### Genome assembly

Guppies for the reference genome assembly were descendants of those collected from a high predation population in the Guanapo River (Twin bridges) from Northern Trinidad (PS 91100 77800). Guppies were inbred through brother-sister mating for 15 generations (Künstner et al. 2016).

From a pool of three full-sib males after 15 inbreeding generations, we generated 50.23 Gb long reads on the Pacific Biosciences RSII platform (read N50 13 kb). Previously available Illumina data (Künstner et al. 2016) was used for polishing. To create a Hi-C library, sperm cells (1.5 × 10^7^) were isolated from a single inbred Guanapo male. Haploid sperm nuclei were approximately 90% as monitored by DAPI staining after lysis. The Hi-C library was prepared as described by Liu and colleagues (2017).

Raw long read data was assembled into contigs with Falcon (version falcon-kit 1.1.1). The resulting primary and alternative contigs were internally phased based on heterozygous SNPs using Falcon-Unzip (version falcon-unzip 1.1.1; parameters default). Phased primaries and alternatives were polished using Quiver (ConsensusCore version 1.02; parameters default). The assembly was phased using Falcon-Phase (commit c86f05e; parameters min_aln_len=5000, iter=100000000, others default). Phase 0 was considered as the primary phase and used for all downstream analysis unless otherwise stated. Completeness of the assembly was conducted by investigating BUSCO scores (Seppey et al. 2019) with BUSCO v.4.0.2, using Actinopterygii as the core ortholog group.

We used slightly updated versions of previous genetic linkage maps ((Tripathi et al. 2009a), GM1; (Künstner et al. 2016); GM2). Both maps used the same cross; GM1 included 776 SNP markers in 896 individuals and GM2 7,509 RAD-seq markers in 184 individuals. We used the maps iteratively, using the more robust GM1 to place most of the contigs, followed by GM2. Finally, we used the genetic map in Bergero et al. (2019), which contained 12 markers to refine the assembly of LG12 (Figure S3, Figure S4).

Our raw assembly consisted of two phases (phase 0 and phase 1). Phase 0 was 751,621,276 bp and phase 1 size was slightly shorter (751,559,625 bp), but there were no major differences in mapping efficiency between the phases (Table S2). Therefore, we used phase 0 for our analysis. We included sequences present in phase 1, but not phase 0, as additional contigs (termed decoys), in our mapping pipeline. The 38 decoys were short sequences totalling 120,900 bp. Note that assignment of contigs to phase 0 or phase 1 is arbitrary, and therefore there is not a simple correspondence between one of the phases and either the X or Y chromosome (see description of mapping decoys in the supplemental materials and Table S4). Differences between the phases were detected by aligning the phases using MUMmer 4 (Kurtz et al. 2004) and SNPs and SVs were called using NucDiff (Khelik et al. 2017).

### Genome annotation

Genome annotation proceeded using BRAKER2 v2.1.1 ((Hoff et al. 2016) https://github.com/Gaius-Augustus/BRAKER), using a combination of GeneMark-EX (Lomsadze et al. 2014) and AUGUSTUS (Stanke et al. 2008; Stanke et al. 2006) using NCBI BLAST (Altshul et al. 1990; Camacho et al. 2009). BRAKER2 was trained using two RNA-seq datasets: one data set was paired-end data from Sharma and colleagues (2014), with tissue from males and females, including whole embryos and multiple adult tissues. The data were accessed via NCBI, Study Accession ID: SRP033586. Illumina sequencing adaptors were removed and reads were quality trimmed to a minimum Phred base quality of 20 as well as discarding broken read pairs, using BBDUK2 (BBMap v34.86, Bushnell http://sourceforge.net/projects/bbmap/); The other data set was single-end RNAseq data from gonad tissue of the ovary of four females, and testes of four males.

RNA-seq reads were mapped to the new reference genome using HISAT2 v2.1.0 (Kim et al. 2019). The alignments were converted to bam format and sorted using SAMtools v1.9 (Li et al. 2009). The guppy genome was soft-masked for repeats using Red (Girgis 2015), a *de novo* approach, which uses a k-mer-based hidden markov model (HMM), and therefore does not rely on previously annotated libraries of repeat sequences. The resulting augustus.gtf file was used downstream for annotation refinement. To refine the annotation, we assessed the Ortholog Hit Ratio (OHR; (O’Neil et al. 2010)). This measure is defined for each unigene with a BLAST match to a related dataset, and is computed by dividing the number of bases in the matched region of the contig by the length of the best-matched sequence with OHR ratios close to 1.0 suggesting complete transcript assembly (O’Neil et al. 2010). To assess OHR, the proteome of *X. maculatus* was downloaded from Ensembl, release 96 (X_maculatus-5.0-male, INSDC Assembly GCA_002775205.2, Dec. 2017). CD-HIT clustering was used to collapse down any peptides with >=90% identity using CD-HIT v4.6.1 (Li and Godzik 2006); (Fu et al. 2012), resulting in a *X. maculatus* peptide database of 27,791 for OHR analyses. gffread (https://github.com/gpertea/gffread) was used to extract protein sequences from the guppy augustus.gtf file and used as a query for BLAST against the *X. maculatus* peptides, using an e-value threshold of 0.00001. Any peptides with an OHR >95% with *X. maculatus* were extracted from the peptide file using samtools faidx (Li et al. 2009) and this well-supported peptide subset was functionally annotated using EggNOG v4.5.1 (Huerta-Cepas et al. 2016), using the eggNOG-mapper v1, mapping with DIAMOND (Buchfink et al. 2015). Emapper annotations were filtered at an e-value threshold of 1e-10.

We also annotated the genome using RepeatMasker ver (open-4.0.5) using the default mode with Actinopterygii as the query (Smit et al. 2019).

### PCR-free libraries for SNP recalibration set

Twelve samples were sequenced, with an average of coverage of 32X, using PCR-free Illumina libraries (Table S13). One male and one female from each of 6 populations, from one river from each of the 3 drainages of the Northern range mountains of Trinidad, with high and low predation (HP and LP) sites for each river (the same populations were sampled for the population genomic study described below using low coverage, Table S5).

Read mapping, cleaning, and SNP calling was similar to that used for our low coverage population sample (see below), except that an additional SNP calling step using using Freebayes v1.3.1 (Garrison 2012) was included, skipping regions with high coverage (-g 1000) and using a minimum alternate allele count (-C 4). Freebayes called 12,726,641 SNPs, and GATK called 13,437,087 SNPs. The intersection of the Freebayes and GATK SNPs was used as our recalibration set (resulting in 8,886,248 SNPs).

### Whole genome alignment

Whole genome alignments between our new male-based assembly (MG), the previous female-based assembly (FG), and the closely related platyfish assembly (Schartl et al. 2013) were done with minimap2 genome alignment (-x asm5, asm20). Alignments were visualised in circos (Krzywinski et al. 2009) and AliTV (Ankenbrand et al. 2017).

### Population genomics

A total of 113 samples were sequenced, including 17-20 individuals from each of six populations across Northern Trinidad (Table S5). DNA was extracted from caudal peduncle tissue using a Qiagen DNAeasy kit. Sequencing libraries were prepared with the Illumina TruSEq Nano DNA library kit, with an insert size of roughly 300bp. Libraries were then sequenced on either an Illumina HiSeq 2000 or Illumina Hiseq 3000, with 5-10 individuals per lane. Each individual was sequenced 1-2 times to obtain approximately 10X coverage.

Data were quality control (FastQC; (Andrews 2010)), cleaned (q20) and adaptors removed (cutadapt; (Martin 2011)) using Trim Galore (Krueger 2015). Reads were aligned to the new MG genome assembly using the bwa mem algorithm in bwa v0.7.17 (Li and Durbin 2009). Alignment files were converted to bam, sorted and indexed using samtools (Li et al. 2009). Validity and quality control of alignments were checked using samtools flagstat and samtools idxstats. Variant calling was conducted using GATKv4 (McKenna et al. 2010), following guidelines for best practices. In cases where more than one lane of data existed per sample, each lane was individually aligned to the genome. Addition of read group (RG) information (AddorReplaceReadGroups) and deduplicating were carried out using Picard-tools v2.17.5 (http://broadinstitute.github.io/picard/). All libraries underwent a first found of deduplicating (MarkDuplicates) for library complexity quality control purposes, before aggregating libraries across samples (MergeSAM) and conducting a second round of deduplicating (MarkDuplicates) to remove any potential library preparation biases. Merged, deduplicated alignments were recalibrated against a variant truth-set using variants generated from our recalibration call set generated from high-coverage, PCR-free sequencing data (see above). Mean and standard deviation coverage for each individual were calculated using qualimap v2.2.1 (García-Alcalde et al. 2012; Okonechnikov et al. 2016).

Variants were called within each population using the GATKv4 processing pipeline for germline variant discovery. SNPs and indels were called via local *de novo* assembly of haplotypes using HaplotypeCaller in genomic VCF (gVCF) mode. Multi-sample joint aggregation of gVCFs was carried out using GenotypeGVCFs, resulting in unfiltered VCF files for each individual. VCF files were filtered to include only biallelic SNPs. Further refinement was conducted using vcftools (Danecek et al. 2011): SNPs were filtered to a minimum depth of 5 and a maximum depth of 200, a maximum amount of missing data of 10% per population and a minor allele frequency of 5%. Only SNPs which were called across all six populations were included for analysis.

Principal components analysis was carried out using PLINK (ver 1.90) (Purcell et al. 2007).

### Coverage

Coverage was estimated from bam files using the bamCoverage option of deeptools (Ramírez et al. 2016). The first and last 100 kb of each chromosome was trimmed before estimating coverage with a binSize = 50, smoothLength=75, and an effectiveGenomeSizeof 528,351,853 bp (genome size minus masked and trimmed regions). The ends of chromosomes were trimmed because they often showed high peaks of coverage, thereby distorting the normalised measures across the chromosome. As the end of LG12 has low genetic content and high GC content, we do not think this caused us to miss any informative sites. Coverage estimates were normalised using RPGC (reads per genomic context). Bins with normalised coverage > 4 (four-times the expected median of 1) were filtered. Coverage was averaged within individuals of each sex within populations and converted to bed files with window sizes of 1 kb and 10 kb by taking weighted means.

Sex-biased coverage candidates were assigned on the basis of log_2_(Male coverage:Female coverage) and defined as greater than 0.75 (male-biased) or less than -0.75 (female-biased). These cut-offs were chosen given expected coverage ratio peaks around 0 (equal coverage), 1 (double male coverage) and -1 (double female coverage).

### Population genomics outliers

Population genetics statistics were calculated using the R package PopGenome (Pfeifer et al. 2014). We calculated F_ST_, D_XY_, and differences in nucleotide diversity (π) in 10 kb windows between sexes within each population. Because D_XY_ is associated with within-group diversity, to assess sequence divergence specifically, we calculated net divergence, D_a_, similar to the definition by Nei (1987). D_a_ can detect sequence divergence between the sexes that cannot be explained in terms of differences in diversity within females (due to mutation rate or other differences). D_a_ was calculated within each population by subtracting female π from D_XY_ calculated between the sexes. Outliers were defined within each population as having values greater than the population mean plus 3 standard deviations (akin to a 99% confidence interval).

While F_ST_ and D_a_ are both based on SNP diversity between the sexes, they will detect different signals. F_ST_ between the sexes will detect areas where all males are heterozygous and all females are homozygous, and when averaging across SNPs over large windows, F_ST_ will only detect these differences if variation between males is low (as expected for a selective sweep of one Y haplotype). D_a_ outliers, however, will detect genomic regions where there is an abundance of low frequency, male-specific variants (that are fixed in females), consistent with the existence of many different non-recombining Y haplotypes.

### Y-mer analysis

Given the low mapping rates to our area of interest (see results), we also took a *de novo* assembly approach to identify male-specific regions. We selected 5 males and 5 females from each of our population samples. Quality of paired-end reads was assessed using FastQC (Andrews 2010) both before and after cleaning and adapter removal using cutadapt (Martin 2011), applying a quality control threshold of 20. The cleaned reads for each sex were pooled for each sample, and error-corrected using Lighter v. 1.11 (Song et al. 2014) using a k-mer size of 31, and a genome size of approximately 730 Mb.

K-mers were counted canonically (-C) for each sex and sampled using Jellyfish v.1.12 (Marçais and Kingsford 2011) with no bloom filter. The ‘dump’ command was used to output the k-mers and their counts. The k-mer abundances were formatted into a matrix using joinCounts from the DE-kupl software (Audoux et al. 2017). A master matrix was constructed containing all k-mers with abundances in each sex for all samples. A second, female-filtered matrix was constructed, containing k-mers which had a frequency <= 1 in females and >=1 in males. This matrix was used to calculate the mean and 95% confidence limits for the k-mers in males in each sample and to define cut-off values for male-specific k-mers, or y-mers.

To identify conserved Y-linked regions, we identified y-mers that were called in all males across all low-coverage population samples. Such y-mers were identified as those with frequency above the sample-specific mean for all male samples in our set of populations, but were not found (frequency = 0) in any females. Reads containing these y-mers were selected from the respective population reads using a custom python script (available). Then contigs containing such y-mers were assembled from the paired-end reads from each population using SPAdes v. 3.12.0 (Bankevich et al. 2012) with the --only-assembler and --careful options. Contig assembly statistics were computed using Quast v 4.6.3 (Gurevich et al. 2013). Contigs for each population were aligned to the genome assembly using minimap2 with the default parameters (Li 2018).

### Investigation of regions of interest

We manually curated gene annotations in our candidate sex determining regions by first searching both for BlastN matches within the reference transcriptome of Sharma et al. (2014) to confirm gene boundaries. The reference transcriptome was searched using BlastN, with the contig as the subject, the 74,567 reference transcripts as queries, and an e-value cutoff of 1e-50. Matching transcripts were then used as queries for BlastX searches on Ensembl 96 of *Xiphophorus maculatus, Danio rerio, Xenopus tropicalis, Mus musculus, Homo sapiens*, and *Drosophila melanogaster*, using an e-value cutoff of 1e-05. These BlastX searches were complemented by domain searches using HMMERscan, with the transcripts translated assuming that each was already in the proper reading frame. The intention here was to characterize sequences in the genomic DNA, not to determine whether they are transcriptionally active.

### PCR validation tests

A PCR presence/absence test for the TRIM-35 duplication was designed with primers at 5,015,401 bp and 5,045,101 bp (see Table S14 for primer details). OneTaq Quick-Load® DNA polymerase (New England Biolabs, Ipswich, MA, USA) was used with 10 pmol of each primer to amplify 1 µl gDNA from individual samples, using the following cycling conditions: 94°C for 1 minute, 35 cycles of: 94°C for 30 s, 53°C for 30 s, 72°C for 60 s; 72°C for 5 minutes. Experiments were performed using a VWR Uno96 thermocycler. The PCR products were visualised using agarose gel electrophoresis with a 2% TAE gel containing a 1:10,000 dilution of SYBR® Safe DNA Gel Stain (Invitrogen, Carlsbad, CA, USA).

Real-time quantitative PCR (RT-qPCR) was performed using primers designed to detect copy number variation of the TeximY gene at 5,186,745 bp (see Table S13 for primer details). iTaq Universal SYBR® Green Mastermix (Bio-Rad, Hercules, CA, USA) was used with 10 pmol of each primer in a 2-step amplification protocol consisting of 30 cycles of: 95°C for 20 s, Ta (53°C) for 30 s. The specificity of the reaction was verified by subsequent melt curve analysis. RT-qPCR was also performed using primers for internal control gene *Rpl7*, under the same conditions. *Rpl7* was chosen as the internal control as it was shown to be the most stable, single-copy reference gene and has a similar efficiency at the same optimal annealing temperature as our target amplicon. A threshold crossing (Ct) value in the target amplicon matching that of the *Rpl7* control was therefore taken as the baseline (i.e. a single copy). The Ct value for each sample was normalised to *Rpl7* and the absolute copy number calculated by comparison to the single copy control samples, using the comparative Ct method (2^-ΔΔCt) (Ma and Chung 2014). Experiments were performed using a Bio-Rad CFX96 Thermocycler, and data processing was performed using Bio-Rad CFX Manager software.

## Data availability

Population genomics data are available on ENA: Study: PRJEB10680

PCR-free data are available on ENA: Study PRJEB36450

Genome assembly is available on ENA ID: PRJEB36704; ERP119926

All scripts and pipelines are available on github: https://github.com/bfraser-commits/guppy_genome

## Acknowledgements

We would like to thank Julia Hildebrandt for sample preparation, Oliver Deusch for data support, and Jenna Cocoran for lab support. We also acknowledge high performing computing (HPC) ISCA server at the University of Exeter.

This work was supported by the Max Planck Society, EU Research Council grant (GuppyCon 758382), NERC grant (NE/P013074/1).

## Notes

### Competing Interest Statement

The authors have declared no competing interest.

