## Supplementary materials for "Improved reference genome uncovers novel sex-linked regions in the guppy (*Poecilia reticulata*)"

**Supplemental Materials:**

### Genome assembly:

*Comparison of phases*

We included sequences present in phase1, but not phase0, as additional contigs (termed decoys), in our mapping pipeline. To examine whether phase1 decoys were potentially biasing the downstream population genetic analyses, we examined coverage of reads to decoy contigs using the same coverage analysis as for the phase0 chromosomes, except that decoys were not trimmed, due to their small size. This analysis was performed with individuals from the GH population from which the genome was sequenced. Coverage of reads to phase1 decoy contigs was generally poor, with 18 and 23 decoys of 38 exhibiting normalised coverage below 0.8 in males and females, respectively, versus only 12 and 1 in phase0 in males and females, respectively. Five decoy contigs exhibited potential signs of sex-linkage, with male coverage greater 0.8 and improved on the phase1 decoy contigs compared with their corresponding phase0 regions. These regions were found at several genome locations, including LG10:1,842,142-1,845,153 bp (phase1 decoy M/F coverage ratio = 2.48, phase0 sex coverage ratio = 0.68), LG10:9,666,131-9,669,307 (phase1 decoy M/F coverage ratio = 3.07, phase0 M/F coverage ratio = 0.79), LG12:2,839,037-2,841,982 (phase1 decoy M/F coverage ratio = 2.12, phase 0 M/F coverage ratio = 0.80), LG12:23,456,655-23,459,300 (phase1 decoy M/F coverage ratio = 2.62, phase 0 M/F coverage ratio = 0.63), and LG8:15,930,097-15,933,408 phase1 decoy M/F coverage ratio = 2.57, phase 0 M/F coverage ratio = 0.62). However, when this analysis was repeated with Oropouche individuals, no windows, including those on LG12, showed consistent male bias across both populations.

**Table S1:** Description of the raw and release genome assembly.

| | **raw assembly** | **release assembly** |
|---|---|---|
| Number of contigs | 401 | 415 |
| Total size | 739303373 | 745001348 |
| N50 contigs | 7935619 | 7936040 |
| N90 contigs | 1441918 | 1331878 |
| Length of longest contig | 25604303 | 25604303 |
| Length of shortest contig | 20028 | 20028 |

**Table S2:** A comparison of mapping rates across different versions of the guppy genome; the previously published female genome (FG), phase 0 of the raw assembly, phase 1 of the raw assembly, and the release assembly. For each population sampled, we summarised the mean per sex of percent of paired-reads mapped properly.

| **Population** | **FG genome** | **MG Phase 0 raw assembly** | **MG Phase 1 raw assembly** | **MG release assembly** |
|---|---|---|---|---|
| HPG females | 93.02 | 97.08 | 97.06 | 97.53 |
| HPG males | 90.14 | 95.95 | 95.59 | 96.36 |
| LPG females | 92.84 | 97.00 | 96.97 | 97.45 |
| LPG males | 90.79 | 95.28 | 95.26 | 95.68 |
| HPO females | 91.49 | 95.08 | 95.07 | 95.09 |
| HPO males | 91.01 | 94.90 | 94.89 | 94.70 |
| LPO females | 91.50 | 94.95 | 94.93 | 94.83 |
| LPO males | 91.35 | 94.92 | 94.91 | 94.83 |
| HPM females | 91.33 | 94.70 | 94.68 | 94.68 |
| HPM males | 91.45 | 94.88 | 94.86 | 94.55 |
| LPM females | 91.10 | 94.44 | 94.41 | 94.05 |
| LPM males | 90.95 | 94.52 | 94.50 | 94.60 |

**Table S3**: Repeat annotation for the male genome (MG) as identified by RepeatMasker ver (open-4.0.5).

| | Number of elements | length occupied (bp) | percentage of sequence (%) |
|---|---|---|---|
| Retroelements | 71565 | 27444545 | 3.68 |
| SINEs: | 21546 | 4586421 | 0.61 |
| Penelope | 265 | 63739 | 0.01 |
| LINEs: | 39941 | 16071894 | 2.15 |
| CRE/SLACS | 0 | 0 | 0 |
| L2/CR1/Rex | 19853 | 4926421 | 0.66 |
| R1/LOA/Jockey | 2612 | 910621 | 0.12 |
| R2/R4/NeSL | 2960 | 2345010 | 0.31 |
| RTE/Bov-B | 11803 | 6195573 | 0.83 |
| L1/CIN4 | 2189 | 1477364 | 0.2 |
| LTR elements: | 10078 | 6786230 | 0.91 |
| BEL/Pao | 958 | 1367916 | 0.18 |
| Ty1/Copia | 95 | 181731 | 0.02 |
| Gypsy/DIRS1 | 5391 | 4500220 | 0.6 |
| Retroviral | 738 | 240562 | 0.03 |
| DNA transposons | 161863 | 26827179 | 3.59 |
| hobo-Activator | 34983 | 3942439 | 0.53 |
| Tc1-IS630-Pogo | 104251 | 20565875 | 2.76 |
| En-Spm | 0 | 0 | 0 |
| MuDR-IS905 | 0 | 0 | 0 |
| PiggyBac | 423 | 84459 | 0.01 |
| Tourist/Harbinger | 1368 | 157179 | 0.02 |
| Other (Mirage, P-element, Transib) | 1 | 73 | 0 |
| Rolling-circles | 0 | 0 | 0 |
| Unclassified: | 8754 | 1427650 | 0.19 |
| Total interspersed repeats: | | 55699374 | 7.46 |
| Small RNA: | 3622 | 768365 | 0.1 |
| Satellites: | 3520 | 815424 | 0.11 |
| Simple repeats: | 289506 | 14785307 | 1.98 |
| Low complexity: | 41722 | 2048748 | 0.27 |

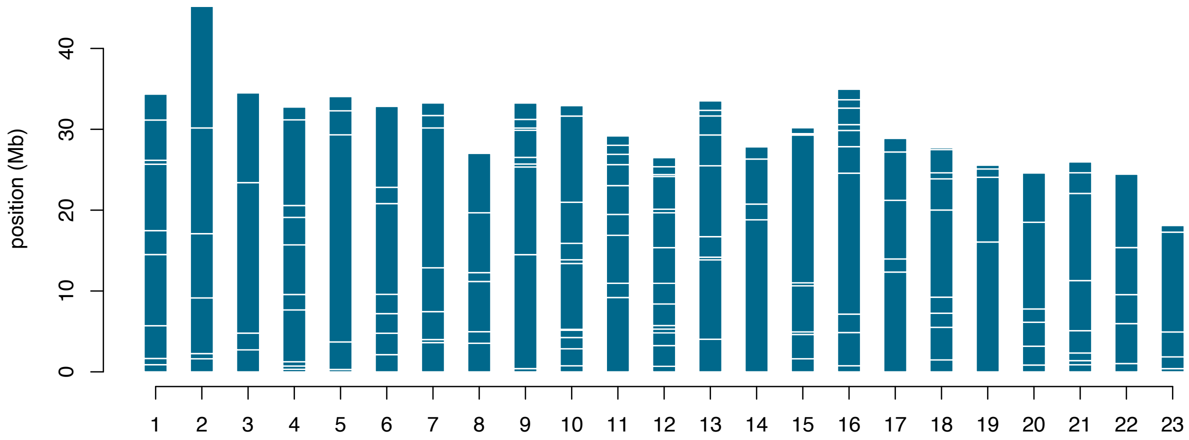

**Figure S1**: Distribution of contigs placed across chromosomes. Contigs are represented as blue boxes.

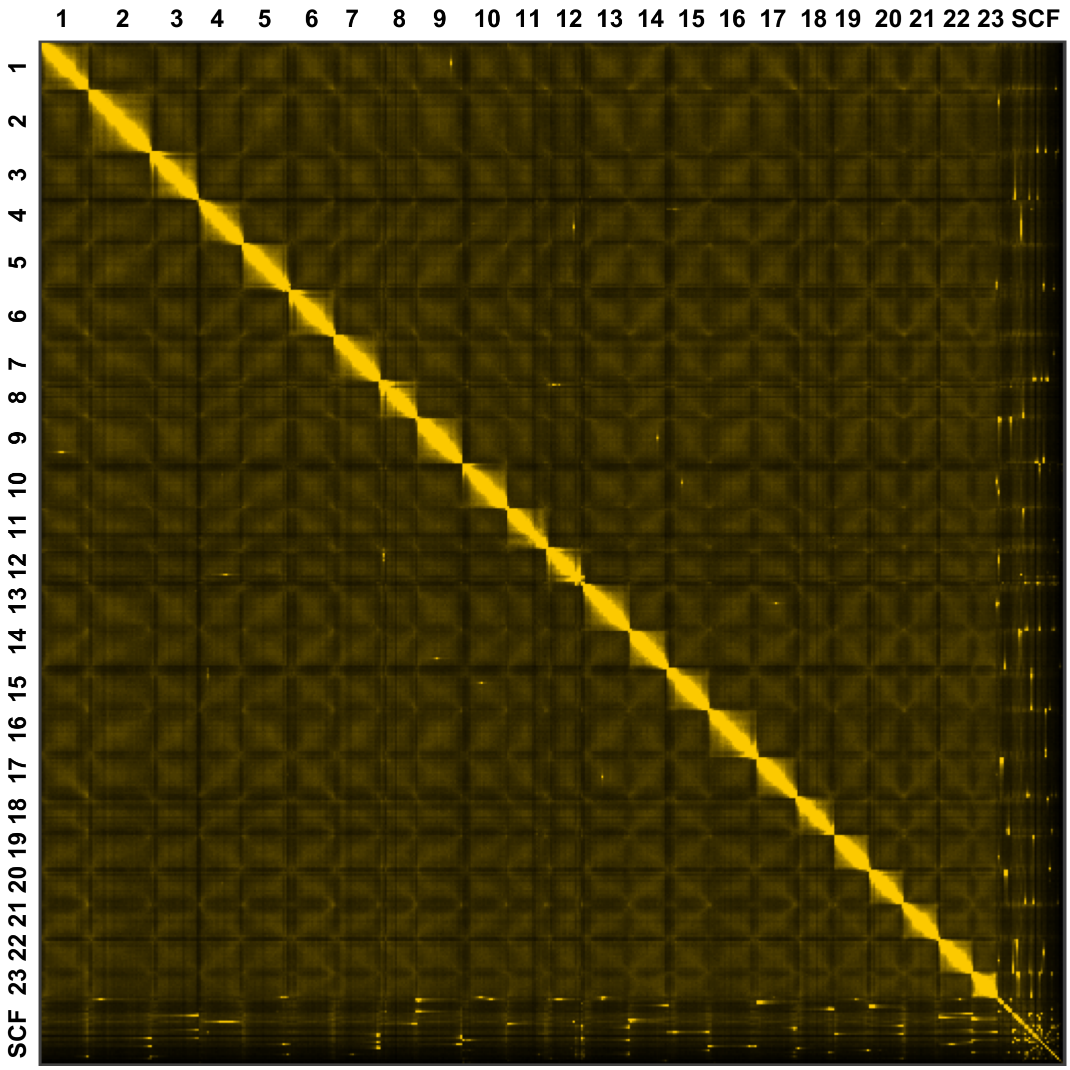

**Figure S2**: Whole genome Hi-C chromosomal contact information, each chromosome is numbered while unplaced scaffolds are placed together.

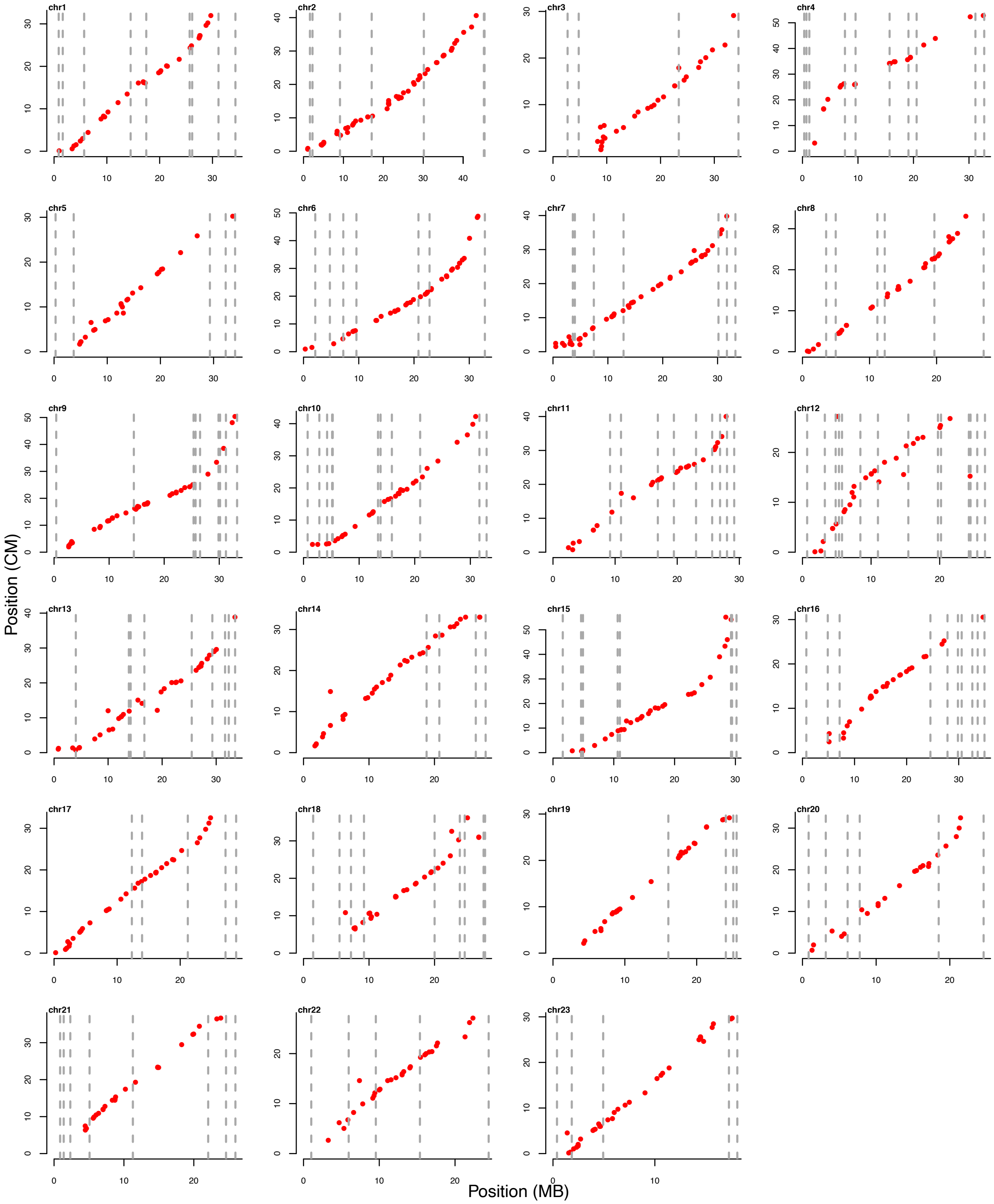

**Figure S3**: Genetic linkage map 1 (GM1), with each marker's position shown on the MG genome in centimorgans (cM). Dashed lines indicate contig boundaries.

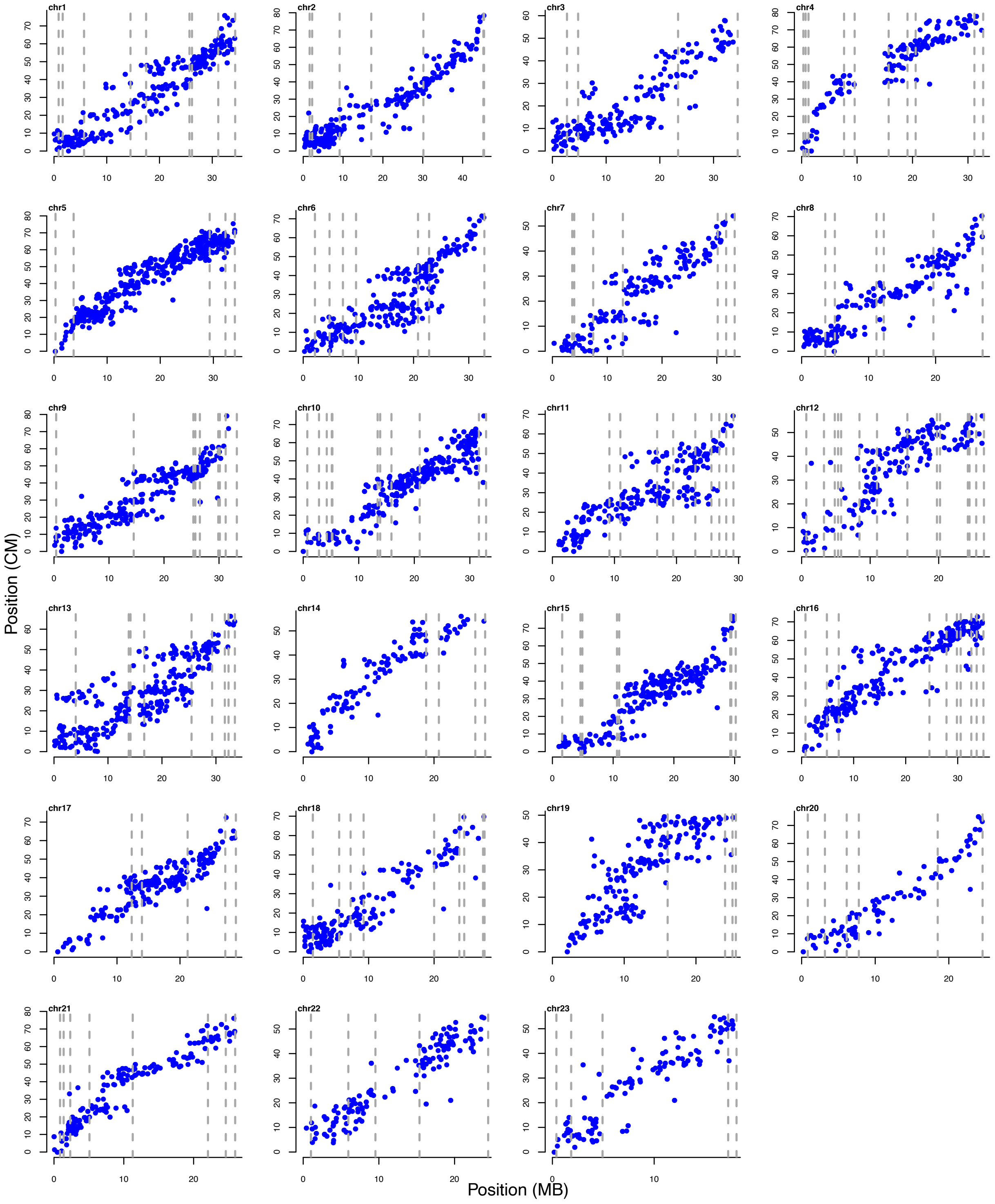

**Figure S4**: Genetic linkage map 2 (GM2), with each marker's position shown on the MG genome in centimorgans (cM). Dashed lines indicate contig boundaries.

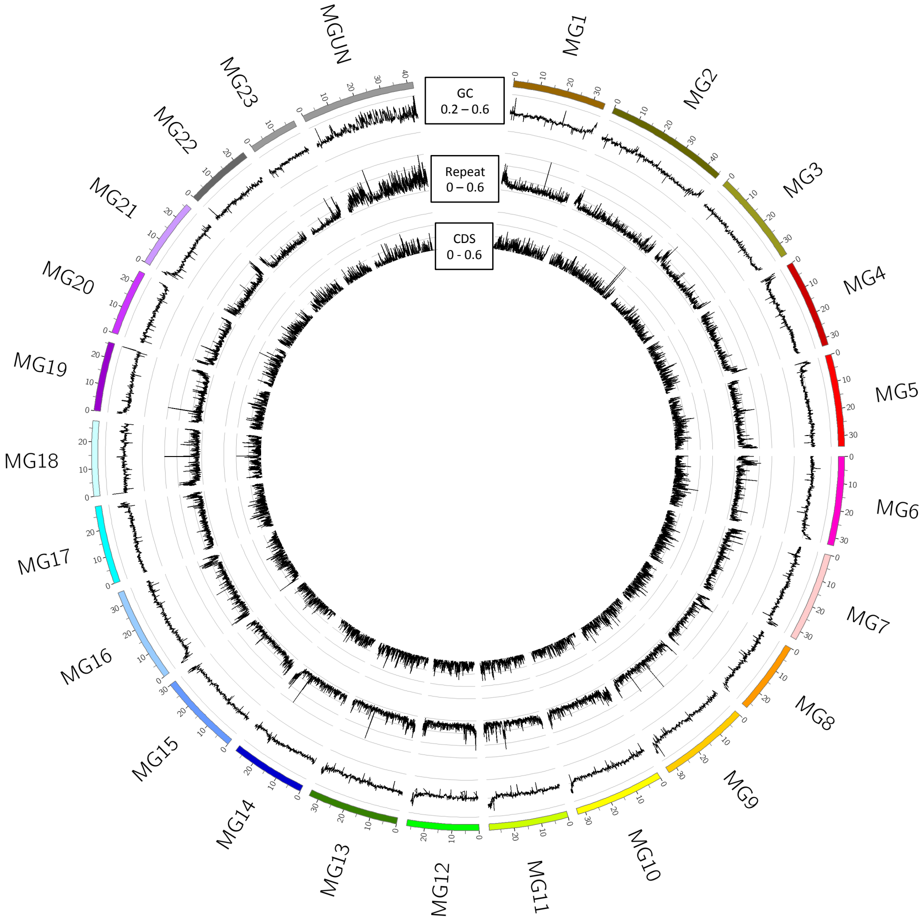

**Figure S5**: Whole-genome distributions of GC content (outermost track), repeat content (middle track), and genic content (innermost track). Each chromosome is shown as well as unplaced contigs as chrUN.

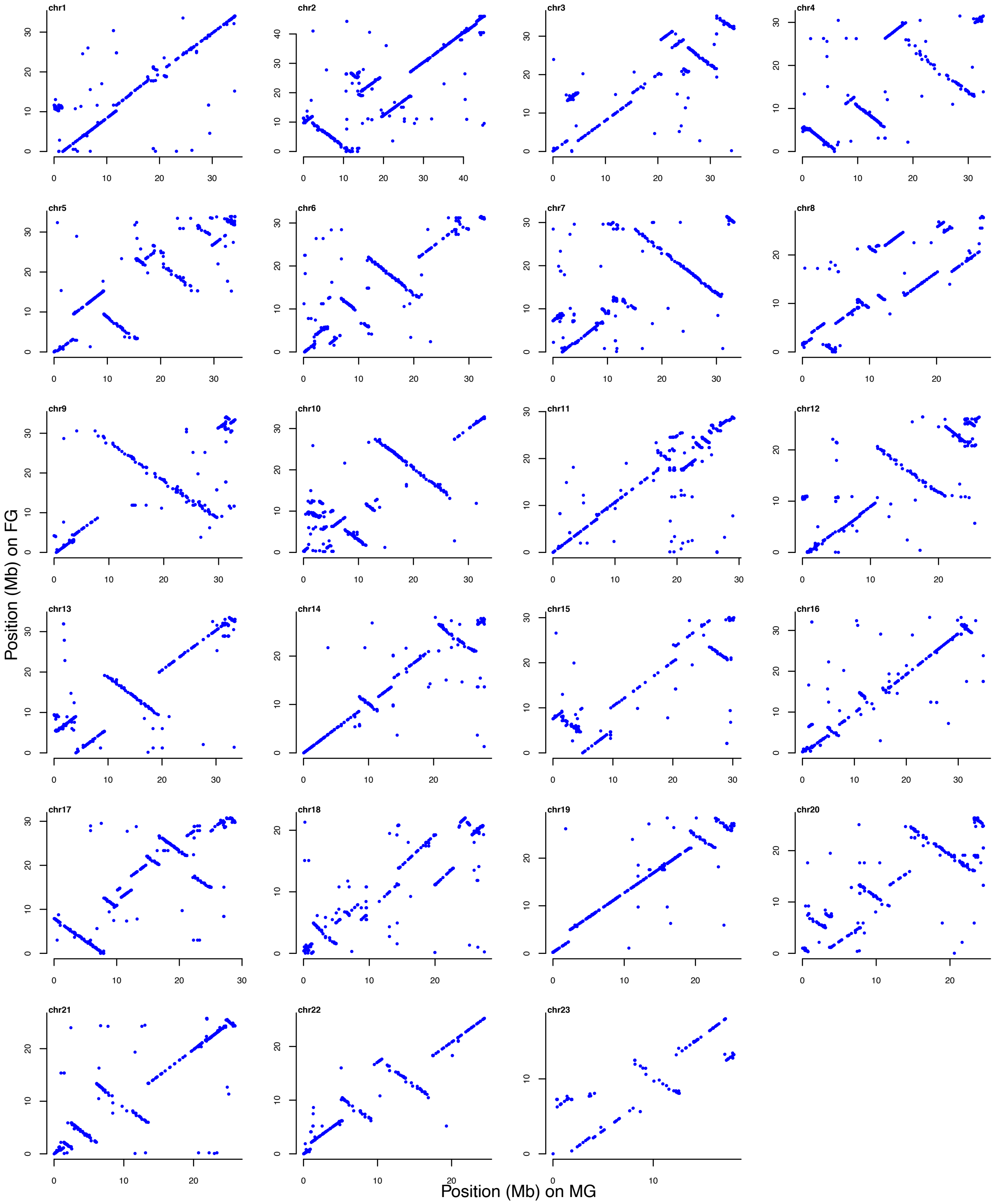

**Figure S6**: Whole-genome alignments between the male genome (MG) and previously published female genome (FG) assemblies for each chromosome. Only alignments in regions with high quality are shown (i.e. mapping quality ≥ 60).

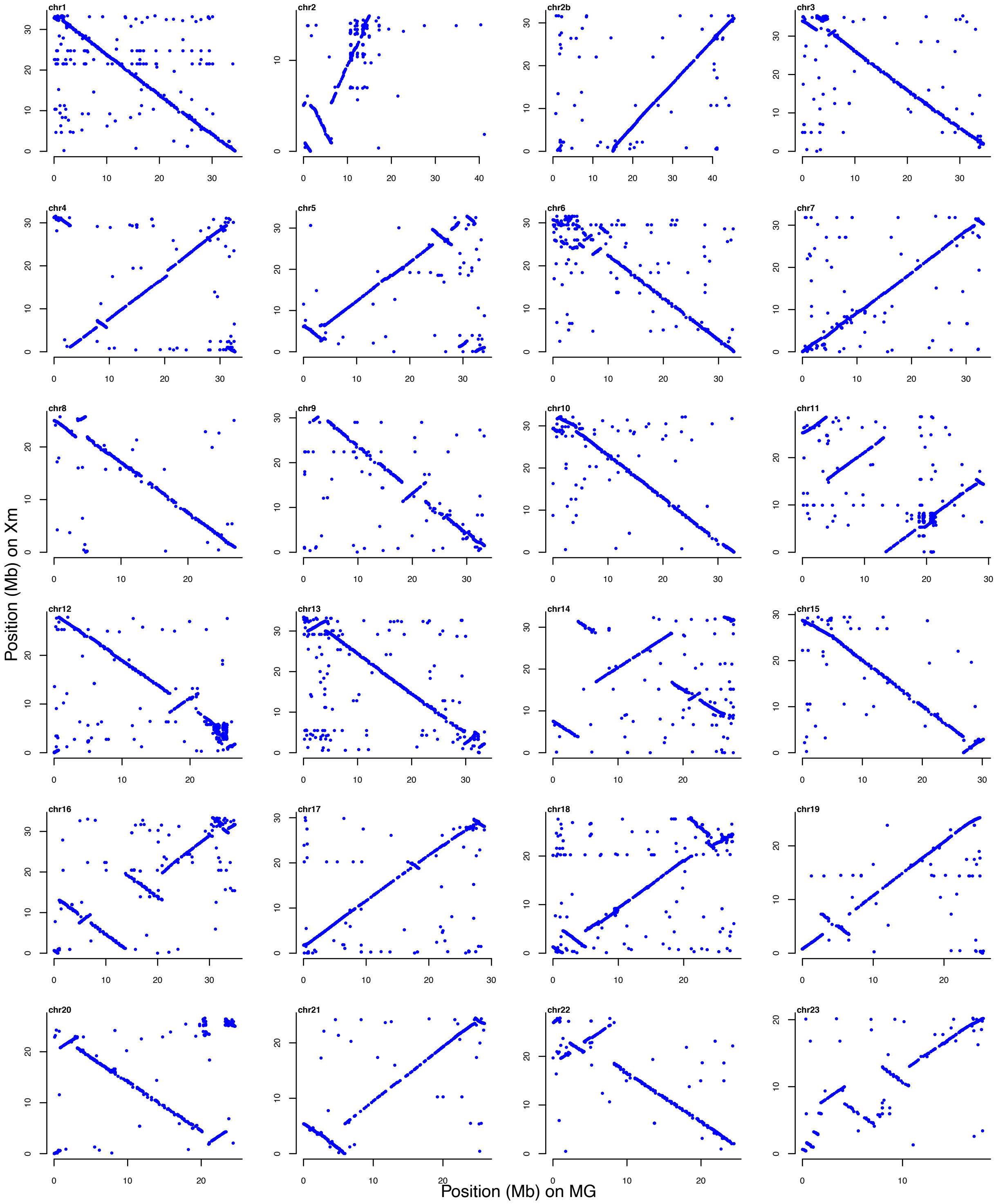

**Figure S7:** Whole-genome alignments between the updated male genome (MG) and *Xiphophorus maculatus* (Xm) assemblies for each chromosome. Only alignments in regions with high quality are shown (i.e. mapping quality ≥ 60).

**Examination of LG12:**

After placing contigs along linkage groups, we revisited the HiC in more detail (figure S8). We rearranged a few contigs so that the ordering was in line with the HiC data. Specifically, we inverted the first two contigs, moved contig V to be adjacent to contig IV, inverted contig XI, and reordered contig XIII and XIV.

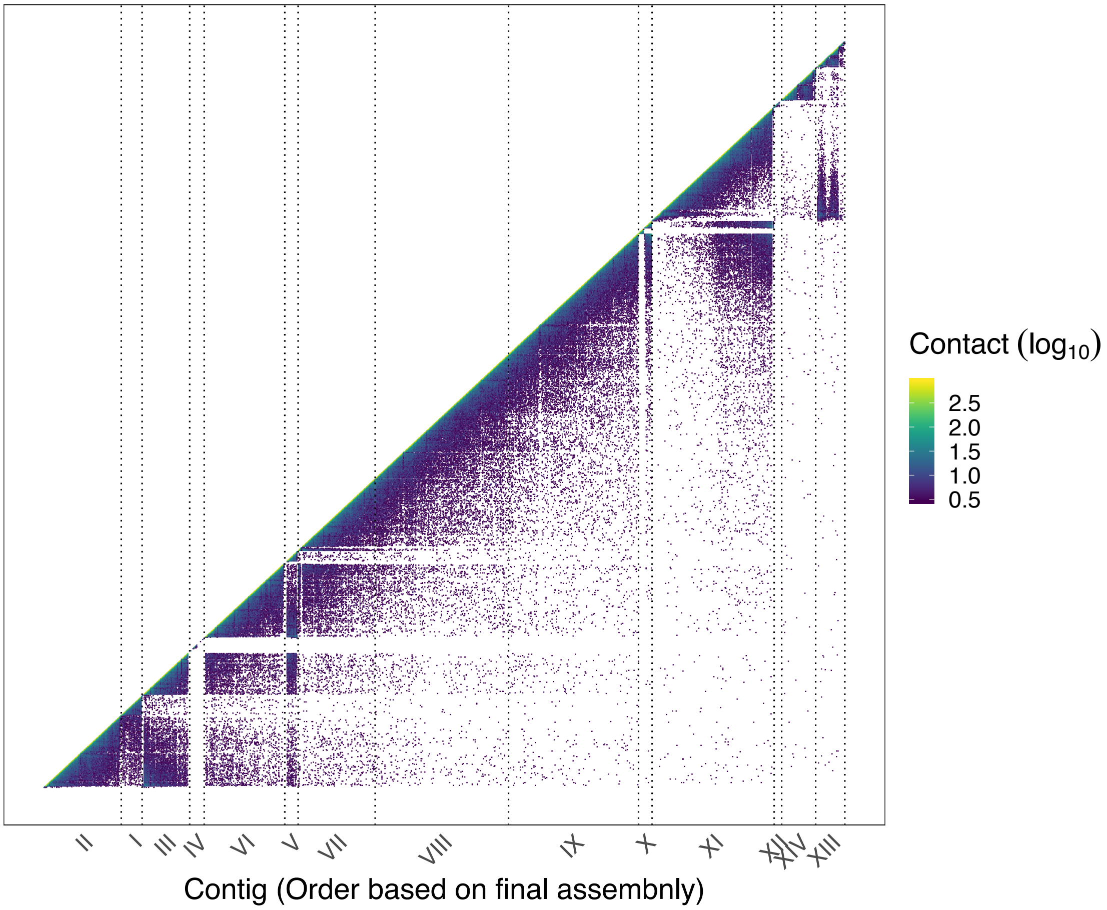

**Figure S8:** Assembly and HiC contact information of LG12 before manual curation, with each contig being delineated with dashed lines and numbered as in main text, i.e. final arrangement.

**Table S4:** Comparison of phase 0 and phase 1 contigs on LG12. Shown are the contig names, corresponding to figure 2, start and end position in the release genome assembly, genetic distance between the phases, and size differences between the phases (phase 0 - phase 1). NA refers to contigs that were changed between the raw and release assembly and therefore do not have corresponding phases.

| Contig ID | Start position | End position | Genetic Distance (SNPs/Contig Length) | Size difference between phases (bp) |
|---|---|---|---|---|
| I | 1 | 682334 | 1.00E-02 | 13667 |
| II | 692334 | 3258872 | 1.22E-02 | -25197 |
| III | 3268872 | 4838750 | 1.11E-02 | 31525 |
| IV | 4848750 | 5321818 | 3.22E-03 | -117381 |
| V | 5331818 | 5765682 | 6.94E-03 | 4464 |
| VI | 5775682 | 8441695 | 9.32E-03 | -62560 |
| VII | 8451695 | 11000781 | 9.62E-03 | 6090 |
| VIII | 11010781 | 15428900 | 9.37E-03 | -11830 |
| IX | 15438900 | 19747371 | 9.56E-03 | 17486 |
| X | 19757371 | 20199082 | *NA* | *NA* |
| XI | 20209082 | 24252561 | 9.79E-03 | 8334 |
| XII | 24262561 | 24501363 | 1.22E-03 | 9 |
| XIII | 24511363 | 25475239 | 9.15E-03 | 7022 |
| XIV | 25485239 | 26605241 | *NA* | *NA* |

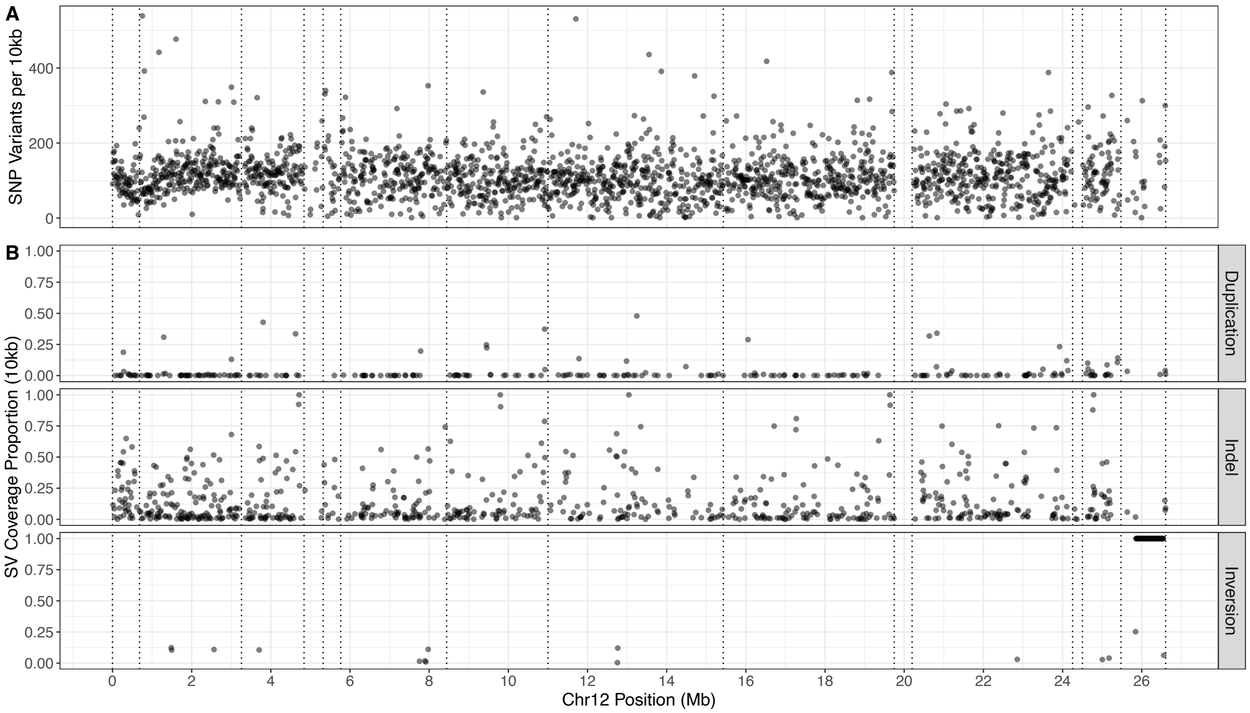

**Figure S9:** Differences between phase 0 and phase 1 across LG12. (A) SNP differences per 10kb, (B) Structural variant (SV) differences summarised per proportion covered by 10kb. Shown are duplications, indels, and inversions. Dashed lines indicate contig boundaries.

**Population genomics**

**Table S5:** Table summarising male and female samples from six wild populations and their mapping statistics to the MG genome. HP denotes high predation, LP denotes low predation populations. IDs of populations given in brackets.

| Population | # of females | Female Samples<br>Mean # reads<br>Mean # mapped paired reads<br>Coverage<br>% paired reads mapped | # of males | Male Samples<br>Mean # reads<br>Mean # mapped paired reads<br>Coverage<br>% paired reads mapped |
|---|---|---|---|---|
| HP Guanapo (GH) | 10 | 77,713,416<br>71,539,143<br>9X<br>97.53% | 9 | 92,662,458<br>86,465,594<br>11X<br>96.36% |
| LP Guanapo (GL) | 8 | 91,316,973<br>83,440,476<br>11X<br>97.45% | 10 | 105,314,493<br>97,036,576<br>13X<br>95.68% |
| HP Oropuche (OH) | 10 | 77,882,229<br>70,619,279<br>11X<br>95.09% | 9 | 106,550,932<br>99,134,444<br>17X<br>94.70% |
| LP Oropuche (OL) | 12 | 86,037,153<br>79,026,947<br>13X<br>94.83% | 8 | 85,358,224<br>78,098,644<br>13X<br>94.83% |
| HP Marrianne (MH) | 10 | 104,765,775<br>95,978,602<br>16X<br>94.68% | 7 | 100,395,645<br>93,492,333<br>16X<br>94.55% |
| LP Marrianne (ML) | 11 | 79,800,263<br>74,532,743<br>14X<br>94.05% | 6 | 105,340,624<br>95,837,332<br>15X<br>94.60% |

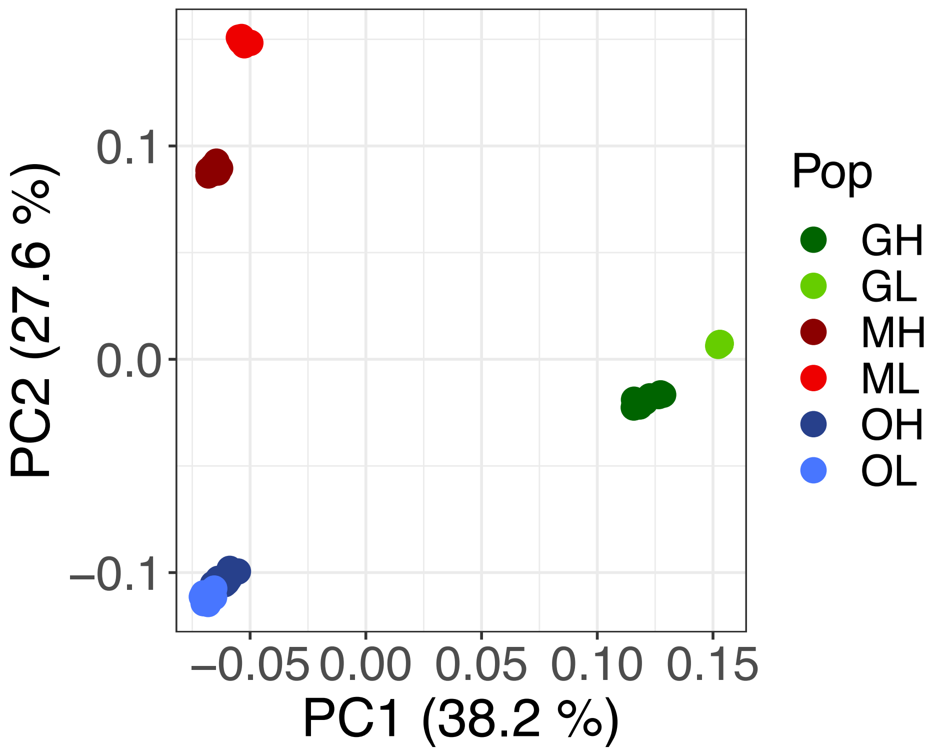

**Figure S10**: Principal component analysis of population samples using whole-genome diversity. Plotted are PC1 (with an eigenvalue of 38.2%) against PC2 (with an eigenvalue of 27.6%), each population is colour coded (see legend).

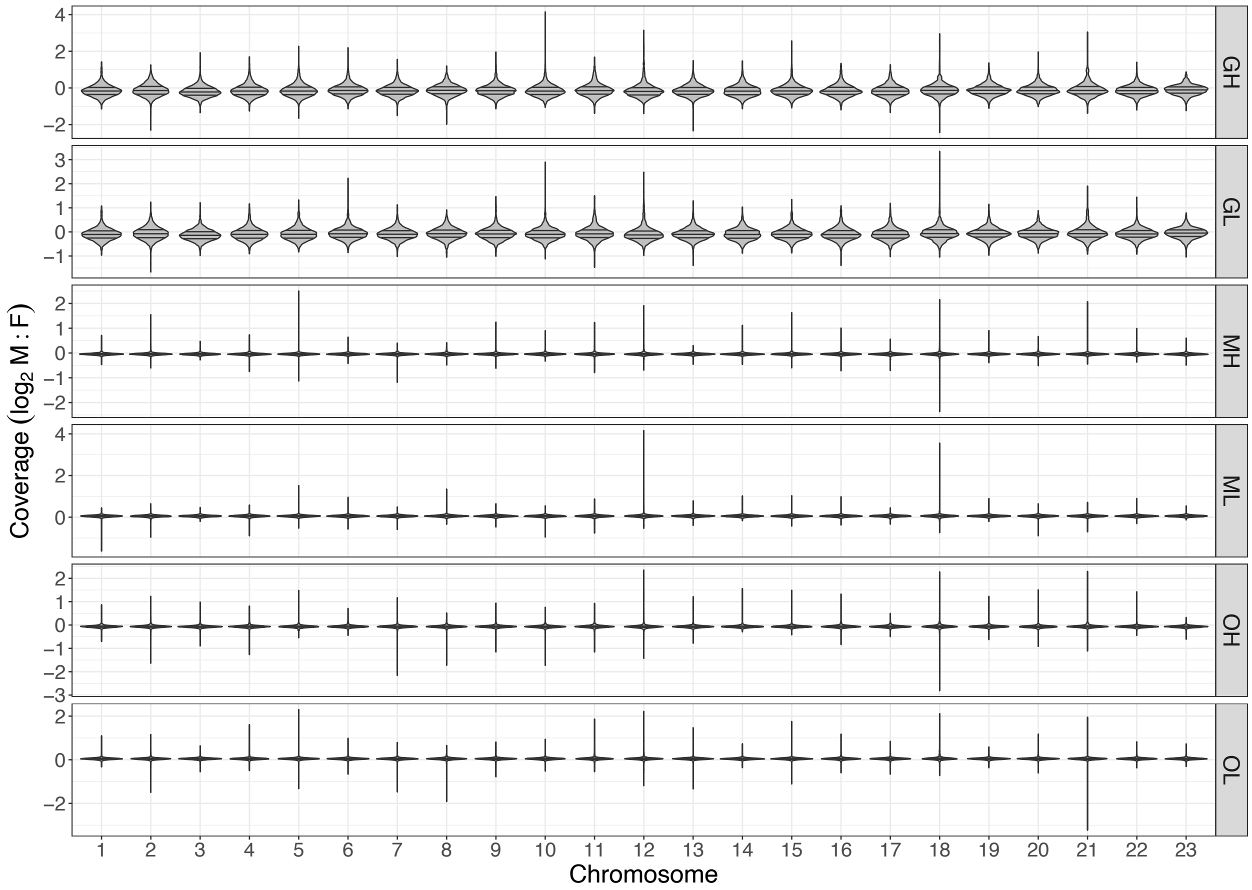

**Figure S11:** Coverage ratio between the sexes (log2[M:F]) across the genome. Shown are violin plots for each chromosome for each population sampled. Population abbreviations can be found in Table S5.

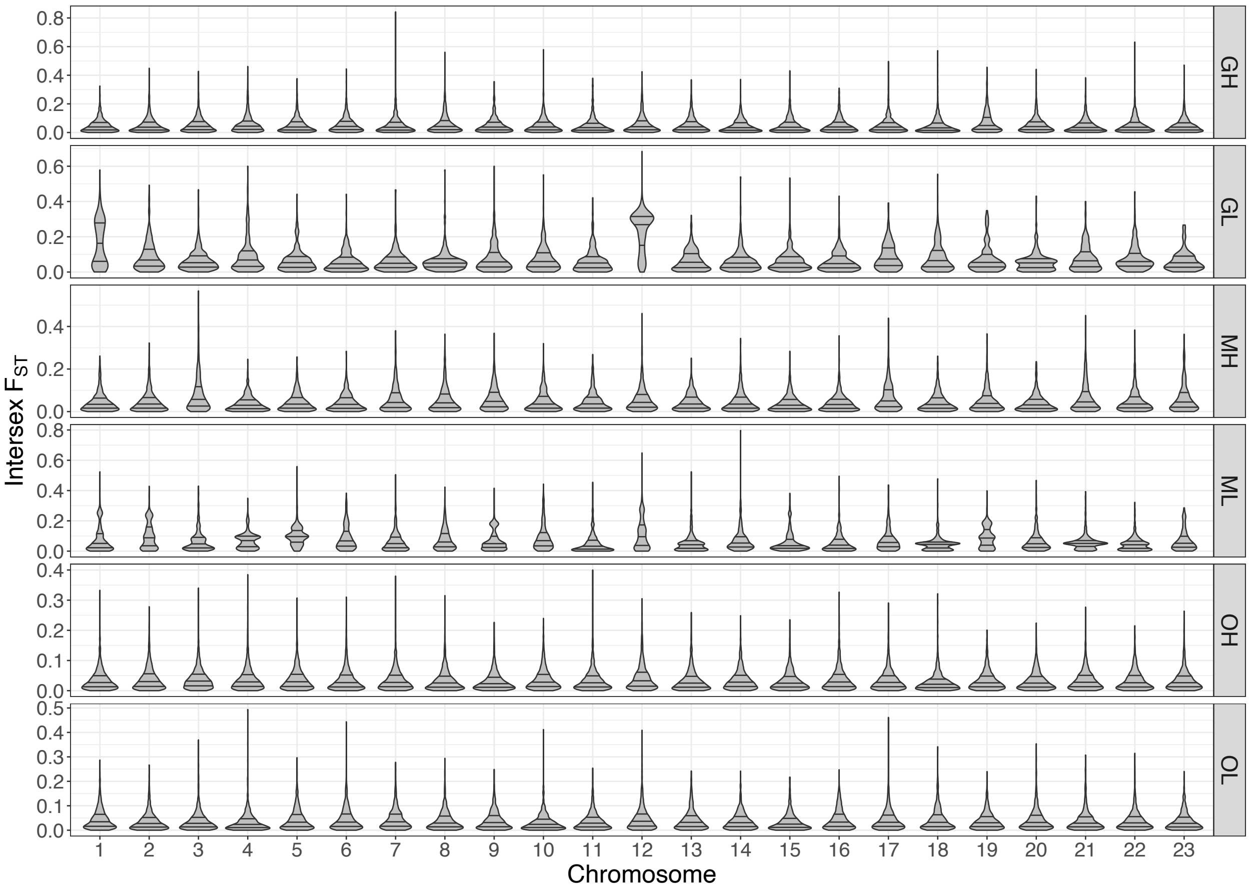

**Figure S12**: $F_{ST}$ between the sexes across the genome. Shown are violin plots for each chromosome for each population sampled. Population abbreviations can be found in Table S5.

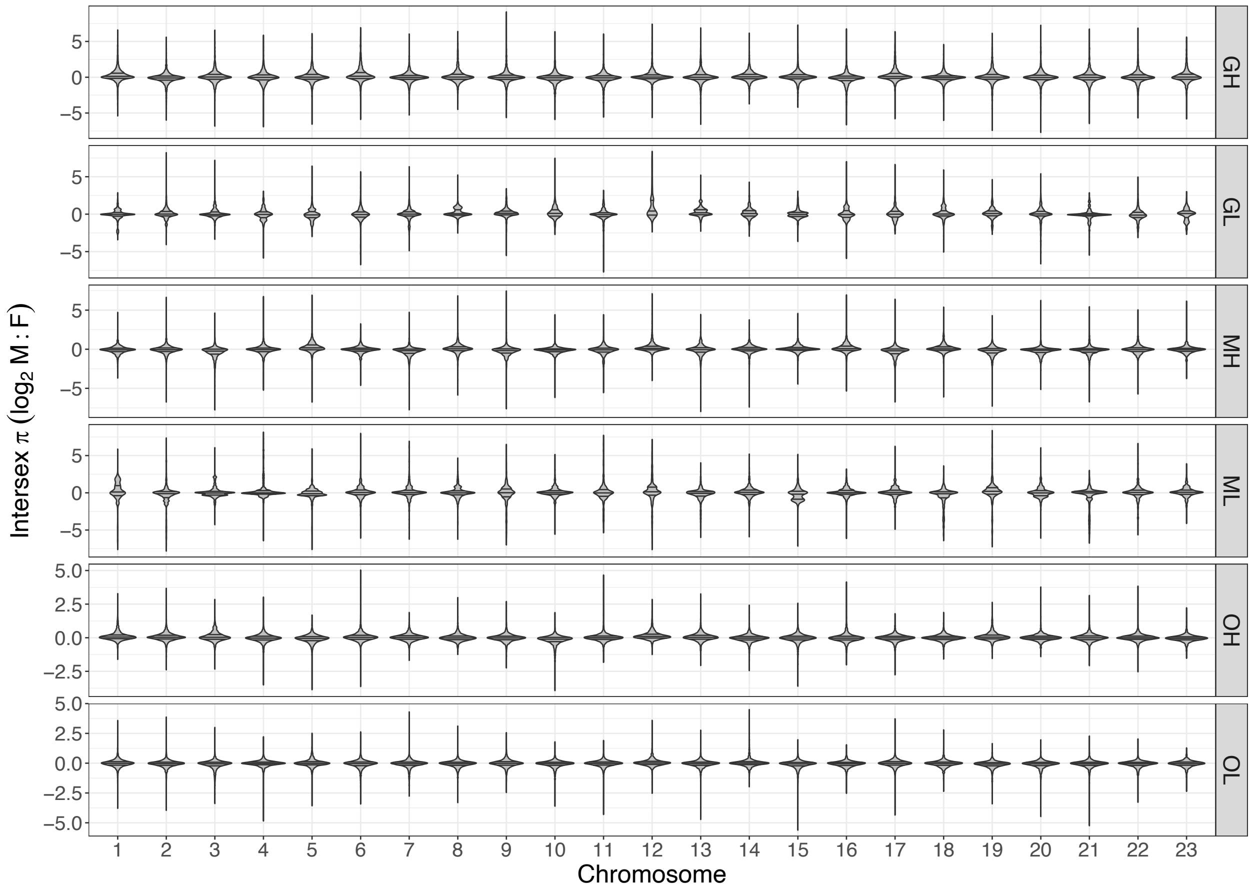

**Figure S13:** Nucleotide diversity (pi) difference between the sexes (log2[M:F]) Shown are violin plots for each chromosome for each population sampled. Population abbreviations can be found in Table S5.

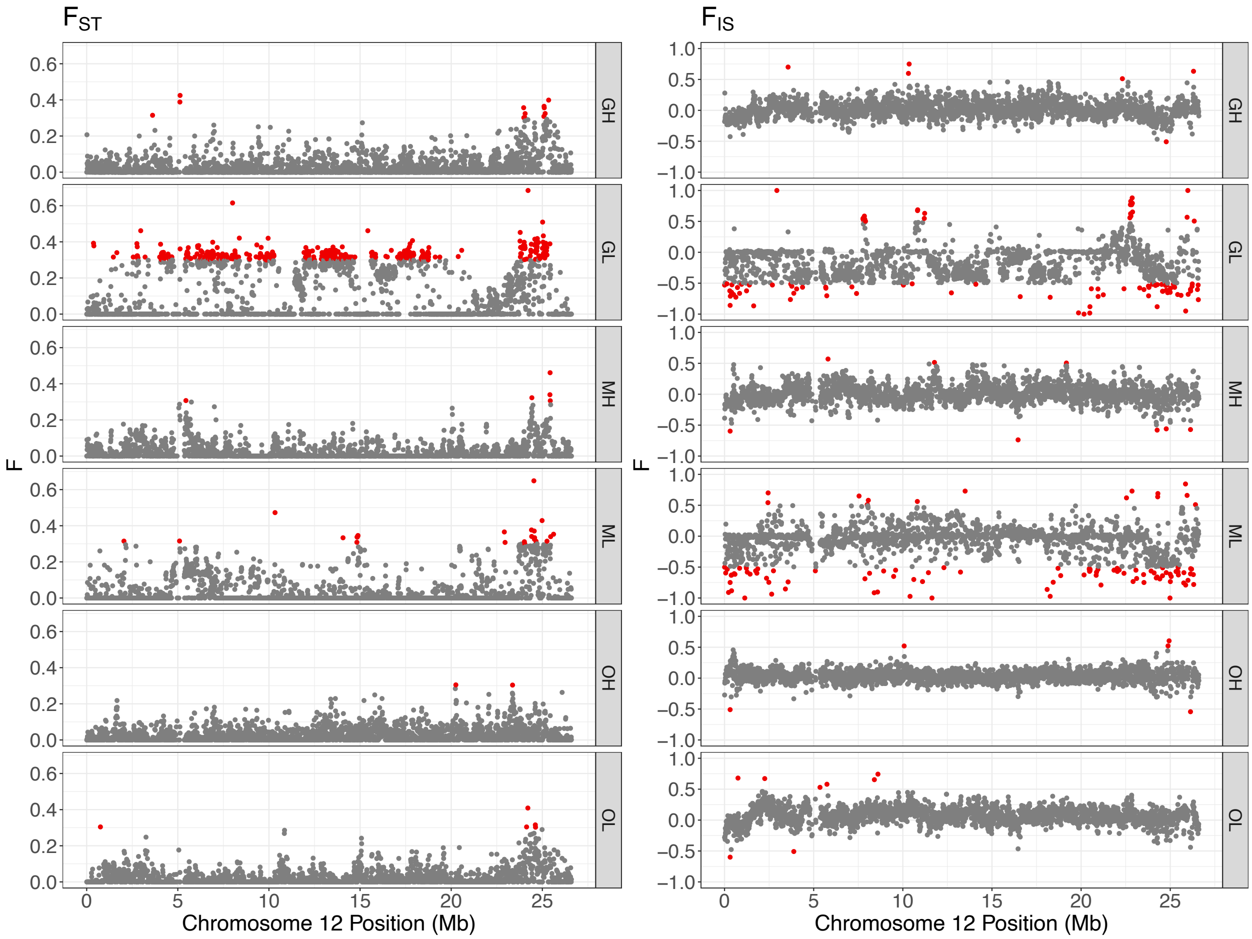

**Figure S14**: $F_{ST}$ between the sexes along LG12 for each population studied, red points denote outliers of above 0.3. $F_{IS}$ between the sexes along LG12 for each population studied, red points denote outliers of above or below 0.5.

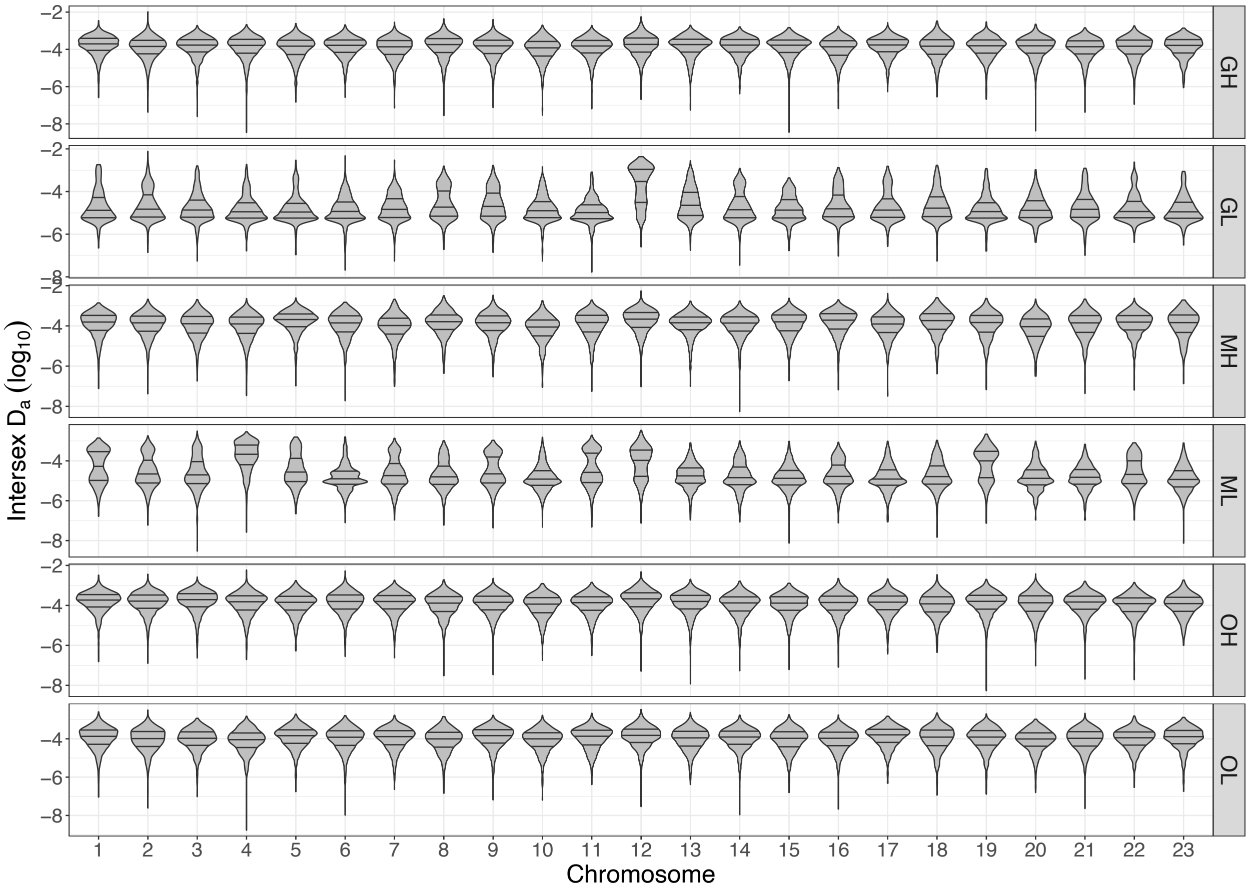

**Figure S15**: Net divergence ($D_a$) along the genome. Shown are violin plots for each chromosome for each population sampled. Population abbreviations can be found in Table S5.

**Table S6:** Overlapping outlier windows of coverage ratio between the sexes. Shown is mean normalised coverage for each sex for each population and the number of populations where that window was identified as an outlier (N_shared). (xlsx file)

**Table S7**: Overlapping outlier windows of $F_{ST}$ between the sexes. Shown are $F_{ST}$ values per population and the number of populations where that window was identified as an outlier (N_shared). (xlsx file)

**Table S8**: Overlapping Net diversity ($D_a$ ) outliers windows. Shown are $D_a$ values per population and the number of populations where that window was identified as an outlier (N_shared). (xlsx file)

**Y-mer results**

After cleaning and error-correction, the average number of reads ranged from 85,938,590 (OL) to 318,421,470 (OH) in males, and the average number of reads in females ranged from 148,775,839 (OH) and 201,770,261 (MH). Between groups, males had a higher number of reads, except in the case of OL. Using the female filtered matrix, the mean number of k-mers ranged from 3 (OL) to 10 (OH) and the upper 95% confidence limits ranged from 12 (OL) to 36 (OH). These values were used as group-specific cut-off values for analysis.

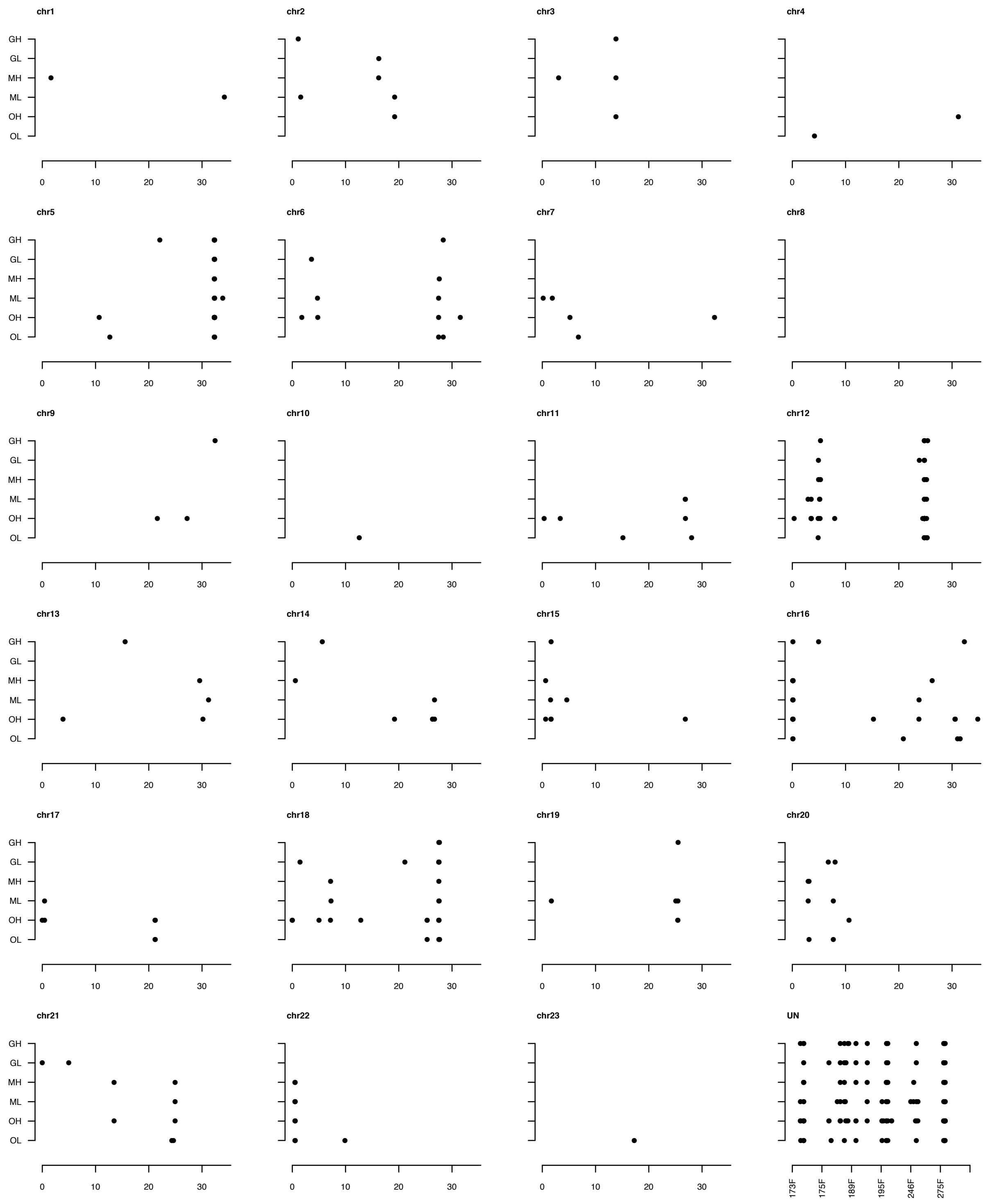

**Figure S16:** Mapping positions of contigs assembled from male specific k-mers, or y-mers, each population is shown on the y-axis across. Panels for each chromosome, and for six unplaced scaffolds that consistently recruited conitgs.

**Table S9**: Summary of y-mers and male specific contigs distributions per population sampled. Shown are number and size of contigs and the linkage groups or unplaced scaffolds they map to.

| | GH | GL | MH | ML | OH | OL |
|---|---|---|---|---|---|---|
| N Y-mer reads | 3546 | 3892 | 3378 | 2710 | 5828 | 2277 |
| N Contigs | 109 | 76 | 92 | 118 | 304 | 95 |
| Total Length of contigs | 37863 | 38489 | 57583 | 77196 | 111551 | 64488 |
| N50 of contigs | 874 | 913 | 1104 | 1041 | 816 | 940 |
| N contigs mapped to genome | 81 | 64 | 80 | 109 | 267 | 85 |
| Length mapped to genome | 34342 | 36173 | 55528 | 74644 | 106566 | 62521 |
| Length on chr12 | 3309 | 3390 | 5065 | 6584 | 11376 | 7430 |
| % | 9.64 | 9.37 | 9.12 | 8.82 | 10.68 | 11.88 |
| Length on chr5 | 3845 | 4098 | 2257 | 8237 | 9760 | 5754 |
| % | 11.20 | 11.33 | 4.06 | 11.04 | 9.16 | 9.20 |
| Length of chr16 | 1661 | 0 | 3176 | 2060 | 2543 | 1829 |
| % | 4.84 | 0.00 | 5.72 | 2.76 | 2.39 | 2.93 |
| Length on chr18 | 680 | 2444 | 1372 | 2664 | 4415 | 2248 |
| % | 1.98 | 6.76 | 2.47 | 3.57 | 4.14 | 3.60 |
| Length of chr22 | 0 | 0 | 1447 | 1853 | 1718 | 4570 |
| % | 0.00 | 0.00 | 2.61 | 2.48 | 1.61 | 7.31 |
| 000173F_0 | 765 | 689 | 2185 | 2603 | 4304 | 3929 |
| % | 2.23 | 1.90 | 3.93 | 3.49 | 4.04 | 6.28 |
| 000175F_0 | 1610 | 1932 | 1495 | 3443 | 3817 | 1840 |
| % | 4.69 | 5.34 | 2.69 | 4.61 | 3.58 | 2.94 |
| 000189F_0 | 676 | 988 | 1806 | 1112 | 1853 | 824 |
| % | 1.97 | 2.73 | 3.25 | 1.49 | 1.74 | 1.32 |
| 000195F_0 | 2972 | 2919 | 6002 | 3354 | 3526 | 5363 |
| % | 8.65 | 8.07 | 10.81 | 4.49 | 3.31 | 8.58 |
| 000246F_0 | 416 | 702 | 247 | 3525 | 1348 | 926 |
| % | 1.21 | 1.94 | 0.44 | 4.72 | 1.26 | 1.48 |
| 000275F_0 | 1637 | 3088 | 5450 | 3881 | 8135 | 4199 |
| % | 4.77 | 8.54 | 9.81 | 5.20 | 7.63 | 6.72 |

**Table S10**: Gene annotations for candidate windows on unplaced scaffolds or autosomes (xlsx file)

**Investigation of candidate sex-determining regions**

Putative protein-coding sequences within contig IV were identified through BlastN searches of the guppy reference transcriptome of Sharma et al (Sharma et al. 2014), using an E-value cutoff of 1e-50. The resulting 273 total matches were reduced to 71 non-redundant matches through an all-against-all BlastN search, retaining only the longest transcript within each cluster of hits. Annotated homologs were identified through BlastX searches of key species on Ensembl 96 (*X. maculatus, D. rerio, X. tropicalis, M. musculus, H. sapiens, D. melanogaster*), using an E-value cutoff of 1e-5. BlastX matches were more frequently observed when searching fish genomes (*X. maculatus* 27/71; *D. rerio* 20/71) than when searching tetrapod or invertebrate genomes (*X. tropicalis* 14/71, *M. musculus* 11/71, *H. sapiens* 12/71, *D. melanogaster* 8/71). Combining results across species, matches were found for 30/71 transcripts; we mostly relied upon annotations from *X. maculatus* (the most closely-related species in the set). Most of the annotated transcripts (19/30) matched known repetitive elements (transposons, retrotransposons, helitrons, retroviruses), while the remainder matched putative “non-repetitive” protein-coding genes. The unmatched transcripts, which tended to be shorter than matched transcripts (IQR of 183–297 nt versus 424–1532 nt), may be non-coding, from highly divergent or de novo protein coding genes, or even artefacts; these were not considered further.

To obtain approximate gene boundaries, the annotated transcripts were mapped onto contig IV using NUCmer (with flags = "-l 10 --maxmatch"). In several cases, multiple transcripts matched common target sequences and mapped to adjacent positions along contig IV, indicating likely split transcripts; boundaries for split transcripts were merged, as needed.

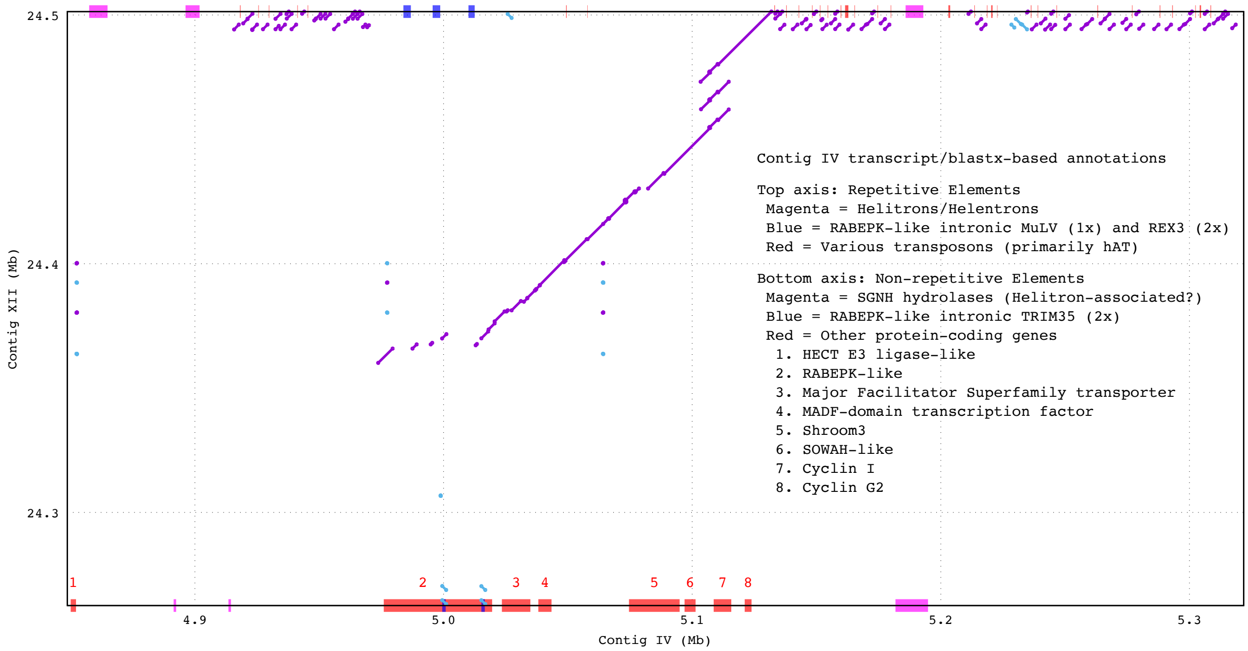

**Figure S17**: Alignment of contig IV (placed at 5 Mb on LG12) and contig XII (placed at 24 Mb on LG12). Alignment dot plots shown in purple (forward alignment) and blue (reverse alignment). Approximate gene boundaries are shown above and below the figure, with annotation names in insert. The gene-rich region (G1 in the main text) is mostly syntenic between the two contigs, with two notable disrupted sections. One, around 5.0 Mb of contig IV, involves a long TRIM35-containing RABEPK intron (see Figure S18). The other, around 5.1 Mb of contig IV, involves a Cyclin I triplication on contig XII.

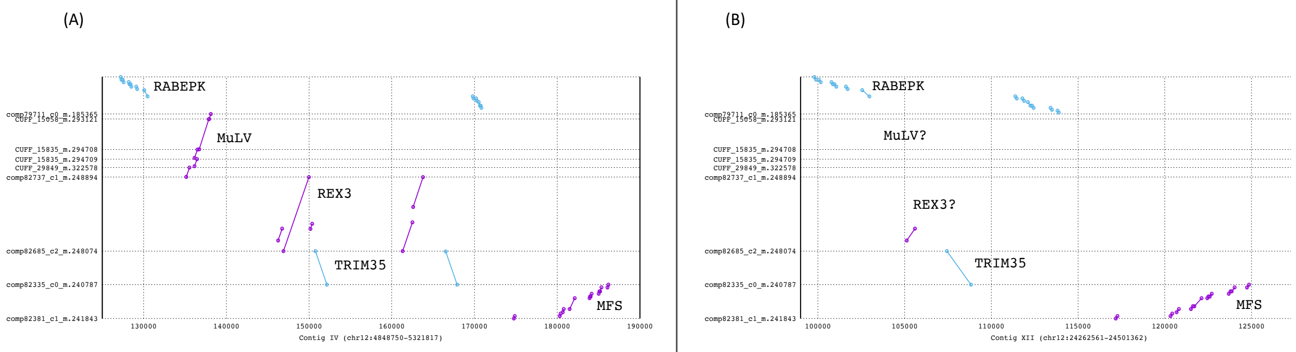

**Figure S18**: Dot-plot alignments of contig IV (a) and contig XII (b) to RABEPK, MuLV, REX3, TRIM35, and MFS transcripts obtained from the reference transcriptome of Sharma et al. (Sharma et al. 2014). Both contigs contain a RABEPK gene followed by a MFS gene. RABEPK possesses a long intron that, on contig IV (a), contains one MuLV retroelement, two REX3 retroelements, and two TRIM35-like genes. The homologous intron on contig XII (b) contains a single TRIM35-like gene.

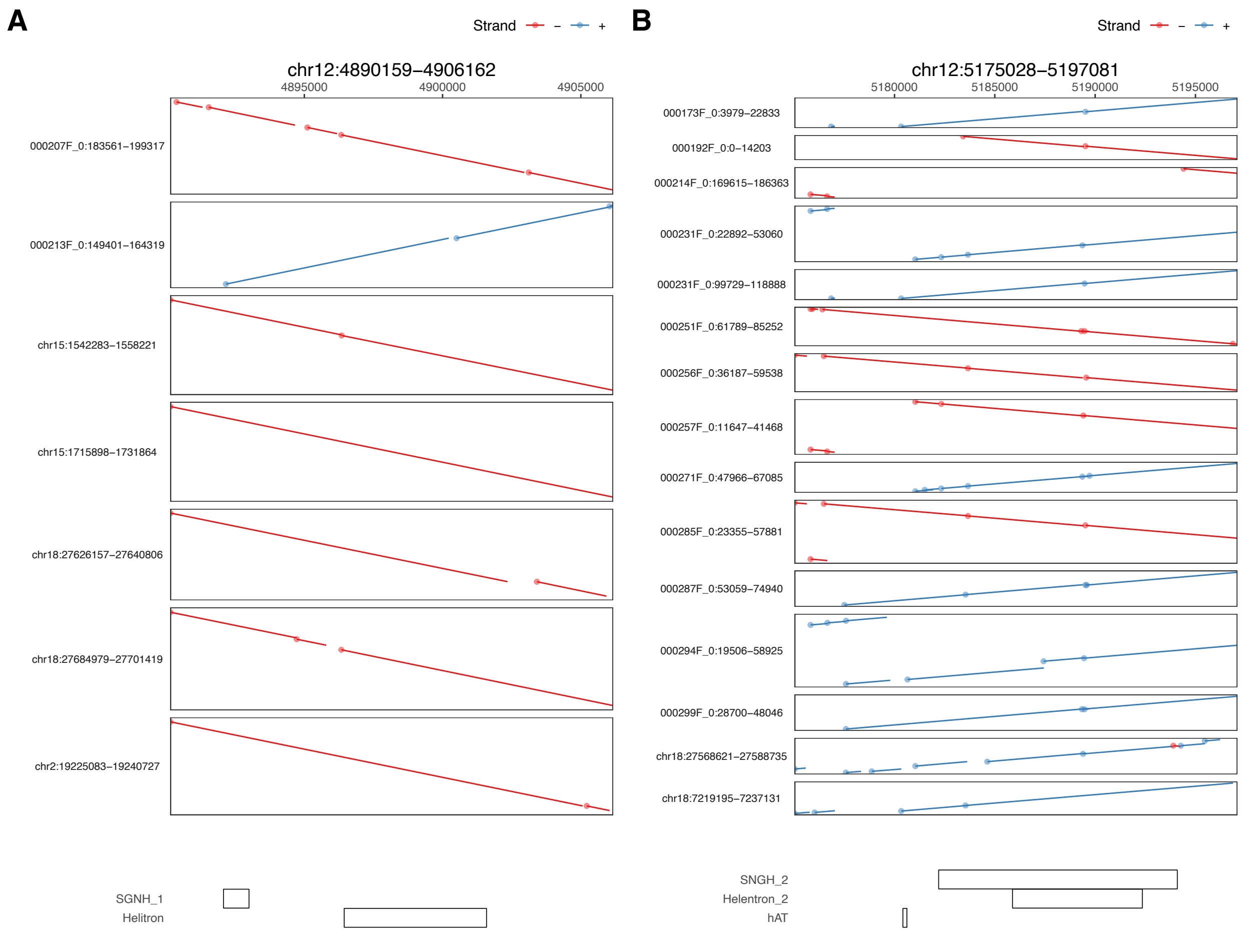

**Figure S19**: Dot-plot alignments of Texim (SGNH helitron genes) regions from LG12 to other chromosomes and unplaced scaffolds. Below the figure is the position of the Texim genes.

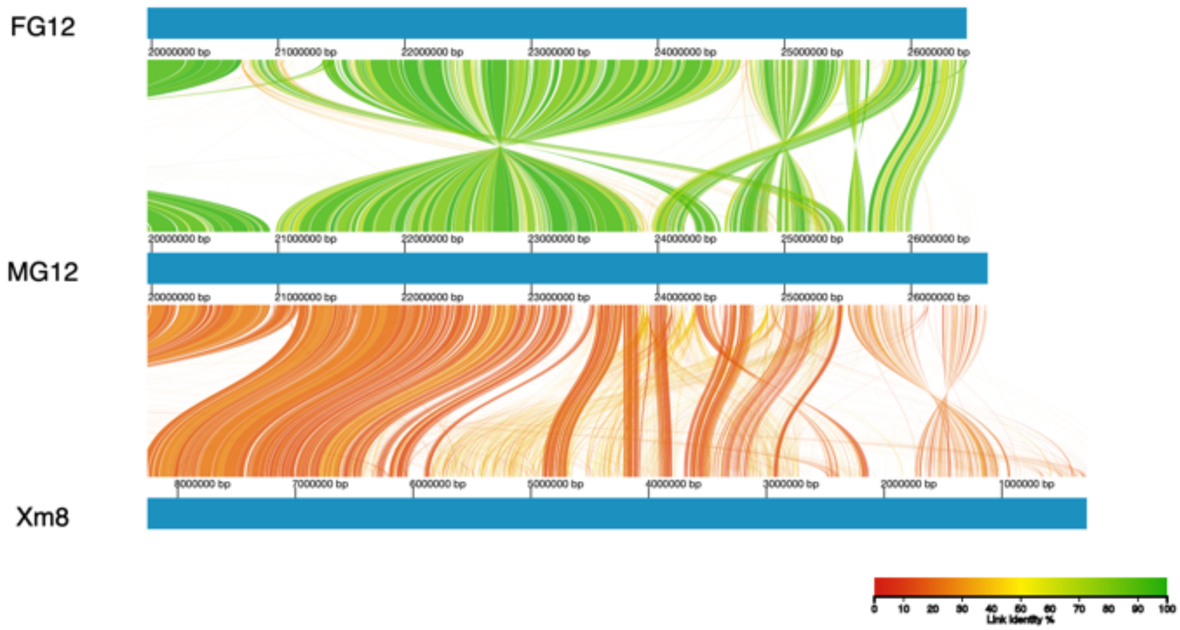

**Figure S20:** Alignment of the distal end of LG12 between the two guppy genome assemblies (female genome assembly: FG and male genome assembly: MG) and chromosome 8 of *Xiphophorus maculatus*. Colour denotes percent similarity of links (see legend).

**Table S11**: PCR test for presence of TRIM-35 insertion summarised per sex per population.

| Populations | Male Present (Total) | Female Present (Total) |
|---|---|---|
| GH | 5 (10) | 5 (10) |
| GL | 7 (10) | 0 (10) |
| OH | 7 (12) | 4 (10) |
| OL | 7 (8) | 1 (12) |
| MH | 9 (10) | 13 (13) |
| ML | 7 (7) | 11 (11) |

**Table S12:** qPCR test for TeximY copy number variation, summarised per sex per population.

| Populations | Male Present mean Log2(ΔΔCt) | Male mean copy number | Female mean Log2(ΔΔCt) | Female mean copy number |
|---|---|---|---|---|
| GH | 2.44 | 2.44 | 0.62 | 1.02 |
| GL | 2.31 | 2.3 | 2.02 | 2.03 |
| OH | 2.18 | 2.19 | 1.74 | 1.53 |
| OL | 1.89 | 1.89 | 1.34 | 1.23 |
| MH | 1.66 | 1.73 | 1.14 | 1.18 |
| ML | 1.76 | 1.77 | 0.51 | 1.00 |

**Table S13:** PCR-free dataset used for recalibration of SNPs, shown are read and mapping statistics.

| Sample | # Raw Reads | # Mapped paired reads | % paired reads mapped | Coverage |
|---|---|---|---|---|
| GH male11 | 114,020,460 | 111,248,558 | 97.72 | 34.51X |
| GH female8 | 103,069,026 | 100,424,044 | 97.62 | 31.15X |
| GL male12 | 121,756,540 | 118,390,104 | 97.89 | 36.33X |
| GL female5 | 66,396,712 | 64,536,388 | 97.87 | 19.83X |
| MH female12 | 141,133,250 | 134,769,650 | 95.72 | 42.23X |
| MH male7 | 134,754,056 | 128,767,236 | 96.25 | 39.82X |
| OH female19 | 131,591,146 | 126,510,152 | 96.78 | 39.04X |
| OH male7 | 99,474,858 | 96,144,274 | 96.79 | 30.07X |
| ML female18 | 155,101,150 | 149,454,766 | 97.01 | 45.90X |
| ML male16 | 133,655,092 | 128,376,742 | 96.73 | 39.65X |
| OL female18 | 54,255,004 | 52,271,994 | 96.49 | 16.28X |
| OL male5 | 62,685,636 | 60,545,666 | 96.69 | 18.89X |

**Table S14**: Primer sequences for PCR and qPCR tests of Table S11 and Table S12, respectively.

| Primer Name | Sequence |
|---|---|
| Trim35 insertion F | CTAAAGATTGCCCACTTCCTG |
| Trim35 insertion R | TCCTCTGTCCCAGTCCAGAT |
| TeximY CNV F | AGCAAAACATCCCGGTTATGT |
| TeximY CNV R | GAGCGTGAAATTGGCCTTCT |
| Rpl7 F | TCAGAGGTATCAATGGTGTCCC |
| Rpl7 R | CAGCTTGACAAACACACCGT |
